# Cardiomyocyte cell cycling, maturation, and growth by multinucleation in postnatal swine

**DOI:** 10.1101/773101

**Authors:** Nivedhitha Velayutham, Christina M. Alfieri, Emma J. Agnew, Kyle W. Riggs, R. Scott Baker, Farhan Zafar, Katherine E. Yutzey

## Abstract

**Aims:** Cardiomyocyte (CM) cell cycle arrest, decline of mononucleated-diploid CMs, sarcomeric maturation, and extracellular matrix remodeling are implicated in loss of cardiac regenerative potential in mice after birth. Recent studies show a 3-day neonatal regenerative capacity in pig hearts similar to mice, but postnatal pig CM growth dynamics are unknown. We examined cardiac maturation in postnatal pigs and mice, to determine the relative timing of developmental events underlying heart growth and regenerative potential in large and small mammals.

**Methods and Results:** Left ventricular tissue from White Yorkshire-Landrace pigs at postnatal day (P)0 to 6 months (6mo) was analyzed to span birth, weaning, and adolescence in pigs, compared to similar physiological timepoints in mice. Collagen remodeling increases by P7 in postnatal pigs, but sarcomeric and gap junctional maturation only occur at 2mo. Also, there is no postnatal transition to beta-oxidation metabolism in pig hearts. Mononucleated CMs, predominant at birth, persist to 2mo in swine, with over 50% incidence of mononucleated-diploid CMs at P7-P15. Extensive multinucleation with 4-16 nuclei per CM occurs beyond P30. Pigs also exhibit increased CM length relative to multinucleation, preceding increase in CM width at 2mo-6mo. Further, robust CM mitotic nuclear pHH3 activity and cardiac cell cycle gene expression is apparent in pig left ventricles up to 2mo. By contrast, in mice, these maturational events occur concurrently in the first two postnatal weeks alongside loss of cardiac regenerative capacity.

**Conclusions:** Cardiac maturation occurs over a 6mo postnatal period in pigs, despite a similar early-neonatal heart regenerative window as mice. Postnatal pig CM growth includes increase in CM length alongside multinucleation, with CM cell cycle arrest and loss of mononucleated-diploid CMs occurring at 2mo-6mo. These CM characteristics are important to consider for pig preclinical studies and may offer opportunities to study aspects of heart regeneration unavailable in other models.

## 1. Introduction

Cardiovascular diseases are the leading cause of mortality worldwide, but effective clinical strategies for regenerative repair of the diseased adult human heart remain elusive.^1, 2^ The mouse heart has been extensively utilized as a mammalian model for cardiac disease in laboratory research, but for clinical studies, the use of large mammals, such as pigs, dogs, and sheep, is mandated. Juvenile pigs, in particular, are often used as cardiac preclinical animal models, due to their hemodynamic and anatomic similarity to the human heart.^3^ However, fundamental developmental characteristics of the porcine heart, including time and pattern of cardiomyocyte (CM) nucleation, hypertrophic growth, and mitotic arrest after birth, are still minimally characterized.

In the first postnatal week, rodent CMs undergo cell cycle arrest with diminishing expression of mitotic markers and cell cycle genes.^4, 5^ At the same time, CMs switch from hyperplastic to hypertrophic growth, with concurrent loss of mononucleation and transition to a binucleated state.^6, 7^ Further, sarcomeric maturation and induction of beta-oxidation metabolism also occur in the first two postnatal weeks in mice, along with an increase in extracellular matrix (ECM) rigidity and deposition.^8, 9^ Thus, by 2-3 weeks after birth, the rodent heart is terminally-differentiated and post-mitotic, forming a fibrotic scar after injury and growing primarily by CM hypertrophy.^10^

Much less is known of these postnatal heart maturational events in large mammals, such as swine. Significant diversity can be conjectured across mammalian species in fundamental cardiac characteristics such as CM nucleation and ploidy, timing of metabolic and hypertrophic switches, and loss of cell cycle activity.^11^ For instance, when considering CM nucleation, adult mice have predominantly binucleated CMs with individual diploid nuclei, while adult humans exhibit high degree of nuclear polyploidy in mononucleated CMs.^12, 13^ Across vertebrates, even more variations, such as tetra- and multi-nucleation, are evident.^14, 15^ However, unlike mice, the postnatal timing of these maturational events, such as loss of mononucleated CMs, cardiac cell cycle arrest, multinucleation, and dynamics of hypertrophic growth, as well as their implications for heart regenerative potential after birth, are still not known in large mammalian models.

Mice exhibit a transient cardiac regenerative capacity in the first week after birth; thus manipulation of CM proliferation and other postnatal maturational events have been identified as potential strategies for inducing regenerative repair after injury in the adult mouse heart.^10, 16^ A similar 3-day transient heart regenerative capacity was recently reported in neonatal pigs, with extensive scarring seen when myocardial injury is induced at P7-P15.^17, 18^ However, the comparative timing of postnatal heart developmental events, as well as the key mechanisms preventing cardiac regeneration beyond the neonatal period in mammals, are not fully known. Direct comparison of cardiac maturational dynamics between different mammalian species could thus help establish the timing of cardiac regenerative potential and the mechanisms underlying postnatal mammalian heart regeneration. For preclinical testing of human therapeutics, pigs are often used as a large animal model.^3^ However, there is a lack of knowledge in basic cardiac cell biology and CM growth mechanisms of the heart in swine, particularly in the postnatal period.

Here, we examine heart maturational events in young swine, including the timing of CM cell cycle arrest and hypertrophy, alongside cardiac metabolic and sarcomeric maturation, to draw a framework for postnatal heart development in a large mammal model in comparison to mice. Our results indicate discordant timing of postnatal cardiac maturation and differential growth mechanisms in pig hearts compared to other mammals.

## 2. Methods

An abbreviated methods section with key experimental details is provided below. The expanded methods section can be found in the online supplementary data file. Details on antibodies and reagents, including dilutions, working concentrations, and manufacturer information, are provided in Supplementary Table 1. Primer sequences used for RT-qPCR are listed in Supplementary Table 2.

### 2.1 Pigs

All experiments with animals were performed conforming to NIH Guidelines on Care and Use of Laboratory Animals and all protocols involving animals were approved by the Cincinnati Children’s Hospital Institutional Animal Care and Use Committee (IACUC). A total of n=68 White Yorkshire-Landrace farm pigs were utilized in this study, with n=14 pigs per stage at P0 to P30, n=4 at 2mo, and n=8 at 6mo. Male and female pigs were included in all analyses.

### 2.2 Tissue harvest and processing

After euthanasia, cardiac left ventricular free wall tissue was obtained for analysis. Freshly harvested tissue was washed in 1X phosphate buffered saline (PBS) and placed in 4% paraformaldehyde (PFA) for histology, in 3.7% formaldehyde for CM dissociations, or flash-frozen in liquid nitrogen for mRNA and protein studies.

### 2.3 Histochemical and immunohistochemical staining

Hematoxylin and Eosin (H&E) staining kit, and Masson’s Trichrome kit were utilized according to respective manufacturer’s protocols to visualize cardiac morphology and total collagen in tissue sections respectively. Capillaries were identified by biotinylated Lectin staining with metal DAB concentrate, and circular Lectin-DAB positive structures were counted using Fiji (ImageJ) software to assess vascular density.

### 2.4 Immunofluorescence staining

Collagen Hybridizing Peptide (CHP, conjugated with 5-FAM) staining, carried out according to manufacturer’s protocol, was used to quantify remodeling collagen. Gap junctional maturation was visually assessed by co-localization of Connexin-43 (Cx43) to intercalated discs. Mitotic indices were calculated by counting nuclear Phosphohistone-H3 Ser10 (pHH3) expression in CMs identified by α-actinin (CM sarcomere) or PCM1 (CM nuclei). Desmin was also used to visualize CMs. Vimentin was used to identify non-CM populations in ploidy quantification experiments. Wheat Germ Agglutinin (WGA, Alexa Fluor 647 conjugate or TRITC conjugate) was utilized to stain for cardiac cell membranes. Nuclei were visualized by DAPI or Hoechst 33342. Staining quantification, object counting, and morphometric analyses were performed using NIS Elements and Fiji (ImageJ) software. Information on antibody catalog numbers and dilutions are provided in Supplementary Table 1.

### 2.5 mRNA analysis by RT-qPCR

Ventricular mRNA was extracted using NucleoSpin RNA kit. Following cDNA synthesis, RT-qPCR was performed using SYBR Green. Gene-specific primer sets for pigs and mice, designed with NCBI PrimerBlast, were validated by Sanger sequencing (Supplementary Table 2). Relative gene expression was calculated by comparative *ΔΔCt* method,^19^ with *18S* ribosomal RNA used for normalization. Normalized average expression of P0 was set to 1.0 to calculate fold change after birth.

### 2.6 Cardiomyocyte dissociations

The protocol of Mollova *et al*.^13^ was modified for isolating individual CMs from pig ventricular tissue. Following 3.7% formaldehyde fixation, pig heart pieces were digested in freshly-prepared Collagenase mixture. Eluate was filtered and gently centrifuged (800rpm, 2 minutes) to obtain CM pellets, which were resuspended in 1X PBS. Antibody staining was performed in cell suspensions by repeated pelleting of CMs after each step. Following staining, CM pellets were resuspended in Vectashield Hardset mounting medium and placed onto slides using coverslips, then dried overnight at 4°C before imaging. A modified method from Patterson *et al*.^12^ was used for ploidy assessment of Hoechst-stained CM nuclei.

### 2.7 Statistics

Statistical analyses were performed using GraphPad Prism 8 software, by unpaired One-way ANOVA non-parametric Multiple Comparisons tests, such as Kruskal-Wallis tests with Dunn’s corrections or Brown-Forsythe and Welch tests with Games-Howell corrections. p<0.05 was deemed significant.

## 3. Results

### 3.1 Cardiac growth by weight is significant at 2mo-6mo in pigs

White Yorkshire-Landrace farm pig hearts were collected at P0 (postnatal day 0), P7, P15, P30, 2mo (2 months post-birth) and 6mo (6months post-birth). The ages were chosen to span birth, weaning, and adolescence in pigs (Figure 1A), similar to the period of cardiac terminal maturation in mice (Figure S1A).^20^ Pig hearts double in anatomical length (atria to ventricular apex) between P0 and P30, with growth to nearly 5 times the size at birth by 6mo (Figure 1B-D). Significant increases in both cardiac and total body weights occurred beyond P30 in postnatal pigs (Figure 1E-F). Further, heart weight-to-body weight ratios (HW/BW in g/kg) increased at P7 and 2mo, but were decreased at 6mo due to disproportionate body weight increases at 2-6mo (Figure 1G-H). When left ventricular morphology in pigs was visualized by Hematoxylin & Eosin (H&E) and Wheat Germ Agglutinin (WGA) staining, myocardial organization into discrete myofibers defined by WGA-staining was apparent from birth, with immature CMs having large central nuclei seen up to P15 (Figure 1I-I’). Hypertrophic expansion of CMs in tissue cross-sections was also only observable at 2mo-6mo. Moreover, the number of nuclei per area (nuclear density) in tissue cross-sections, used as a gross measure of cell density, was not significantly reduced until 6mo (Figure 1J). In contrast, myocardial organization in mice is evident by P7 with hypertrophic growth observable by P15 (Figure S1B). Together, these data suggest a cardiac terminal maturational period several months after birth in pigs, whereas in mice these maturational transitions occur within a few weeks after birth.

**Figure 1.**
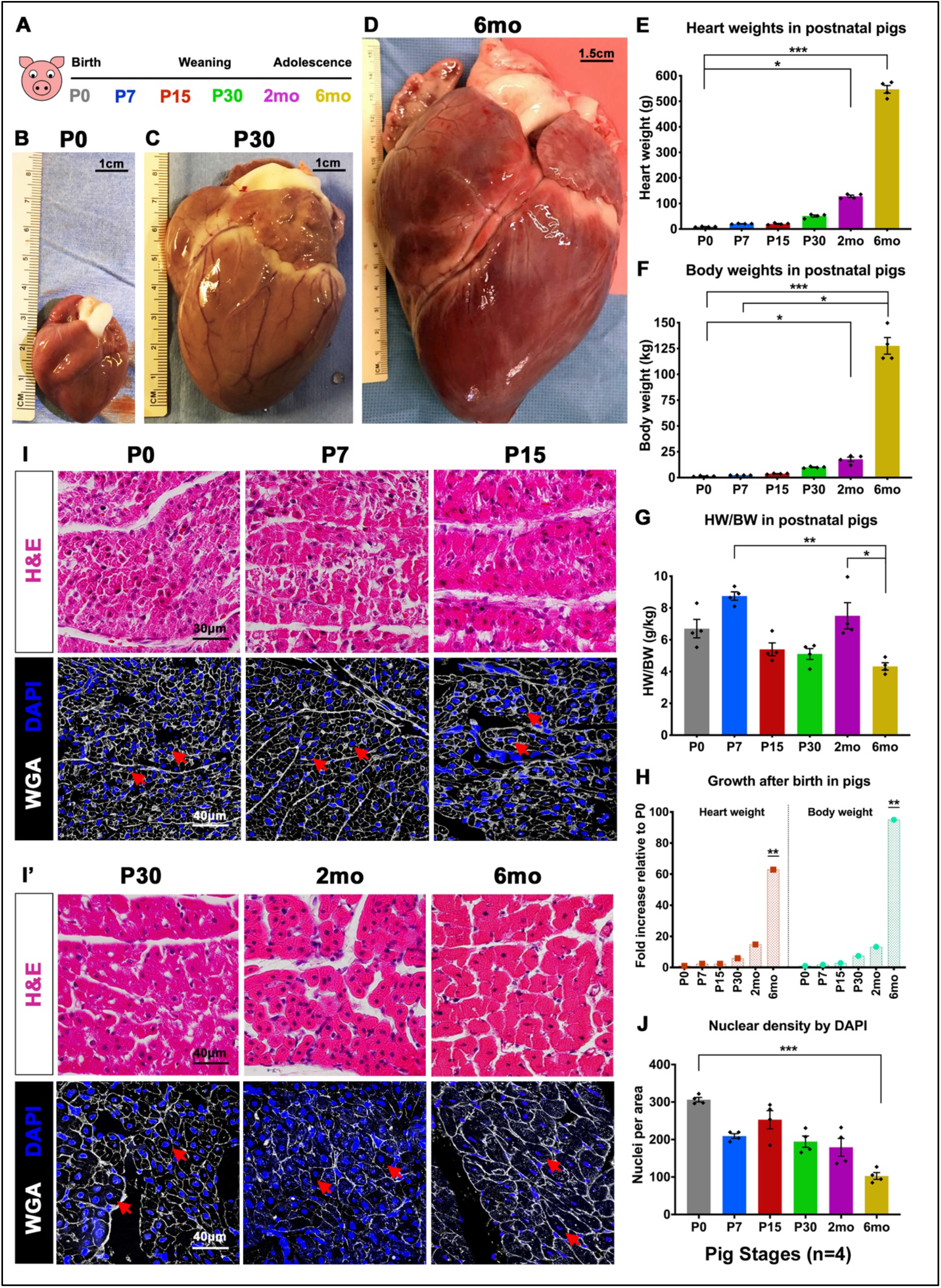
Postnatal pig hearts increase significantly in size beyond P30. **(A)** Schematic showing pig heart stages utilized for study. **(B, C, D)** Representative images of postnatal pig hearts at birth (P0), one- (P30) and six-months (6mo) post-birth, with relative sizes indicated in centimeters (cm). **(E)** Heart weights were measured in grams (g) at P0-6mo in pigs. **(F)** Total body weights were measured in kilograms (kg) at P0-6mo in pigs. **(G)** Ratio of heart weight/body weight (HW/BW in g/kg) was calculated for pigs at P0-6mo. **(H)** The increase in heart and body weights after birth in pigs is shown relative to P0 weights. **(I, I’)** Myocardial histology was analyzed by Hematoxylin & Eosin (H&E) and Wheat Germ Agglutinin (WGA) with DAPI staining of nuclei in postnatal pigs. Red arrows indicate cross-sections of CMs as indicators of cell size expansion. **(J)** The number of nuclei per area (0.05mm^2^) was assessed by DAPI counts in P0-6mo pig left ventricles. Data are mean ± SEM, with *p<0.05, **p<0.01, ***p<0.001 determined by Dunn’s Kruskal-Wallis Multiple Comparisons Tests, in n=4 pigs per stage.

### 3.2 Vascular density does not change after birth, while collagen remodeling continues to 6mo in postnatal pigs

Maturation of the postnatal heart in mice is characterized by increased vascularity and ECM deposition, which are also implicated in myocardial regenerative capacity.^21, 22^ Vascular density was assessed by Lectin-DAB staining in postnatal pigs and mice as an indicator of myocardial vascularization and maturation after birth. Capillaries stained by Lectin-DAB were quantified in pig hearts at P0-6mo, in parallel with P0-P30 mouse hearts. In pigs, cardiac vascular density did not change from birth to 6mo (Figure 2A-B). In mice, however, there is trend towards increasing vascular density in the first week after birth (Figure S2). Therefore, myocardial vascularization is established before birth in swine, contrary to the postnatal vascularization and remodeling observed in rodents at P0-P7.

**Figure 2.**
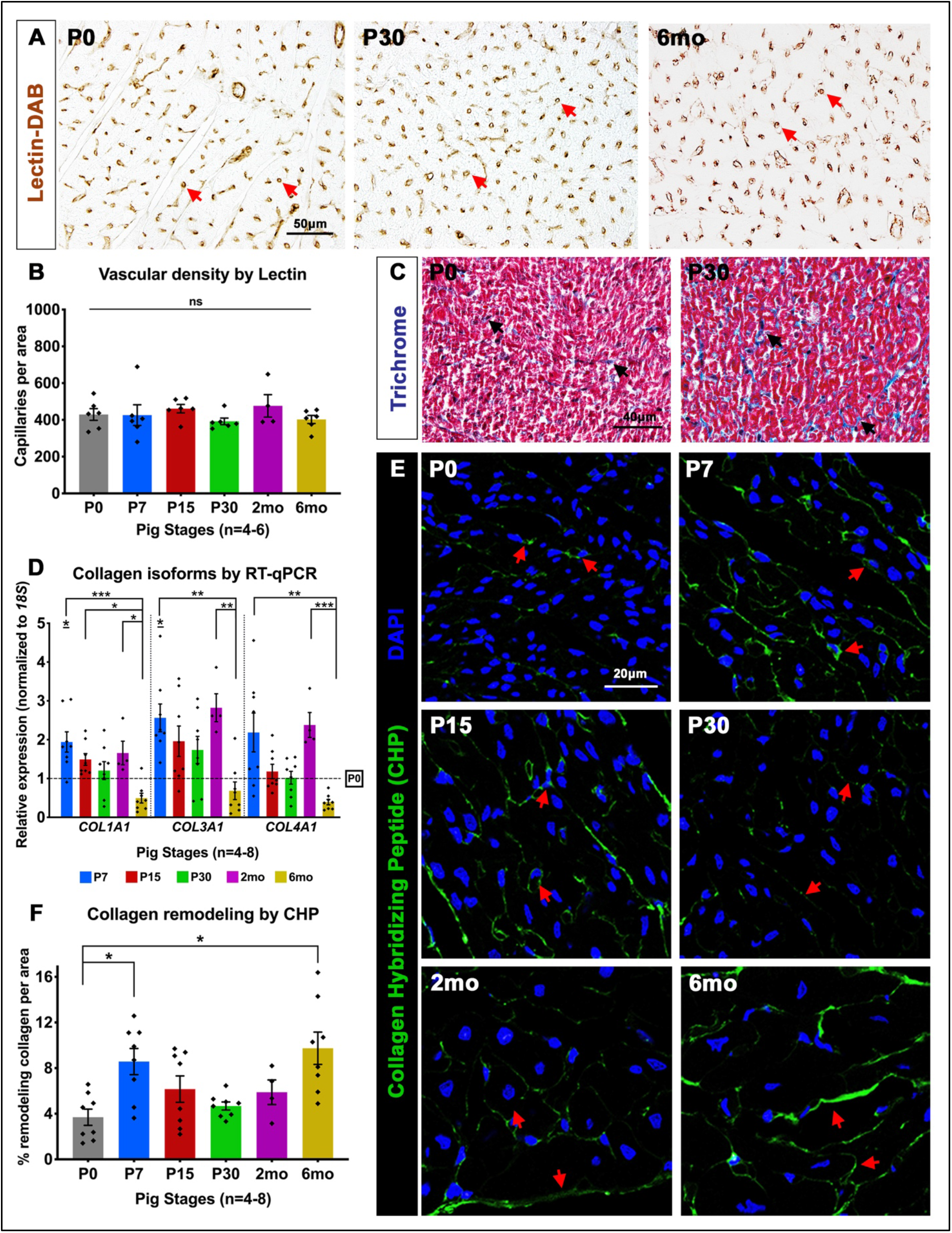
Vascular density does not change after birth while collagen remodeling increases at P7 in postnatal pig hearts. **(A)** Representative images showing vascular capillaries stained by Lectin-DAB in pig hearts. Red arrows indicate circular capillary structures. **(B)** Vascular density was assessed by counting Lectin-DAB stained capillaries per area (0.3mm^2^). **(C)** Representative images of Masson’s Trichrome staining showing fibrillar collagen (black arrows) at P0 and P30. **(D)** RT-qPCR analysis of collagen isoform genes at P7-6mo with fold change indicated relative to P0 in pig left ventricular mRNA. **(E)** Collagen remodeling was assessed by fluorescent Collagen Hybridizing Peptide (CHP) staining (red arrows). **(F)** Remodeling collagen was quantified as the percent area of CHP expression per total area (0.05mm^2^), with significant increase at P7 and 6mo. Data are mean ± SEM, with *p<0.05, **p<0.01, ***p<0.001 determined by Dunn’s Kruskal-Wallis Multiple Comparisons Tests, in n=4-8 pigs per stage (ns=not significant). In D, asterisk(s) with underline indicate significance compared to P0.

Postnatal ECM maturation has also been previously indicated as a key event in heart maturation and CM cell cycle exit within one week after birth in mice.^23^ The timing of ECM maturation in postnatal pigs compared to mice was evaluated by collagen deposition, gene expression, and fiber remodeling. Masson’s Trichrome staining revealed appreciable fibrillar collagen at birth in pig hearts, with an apparent increase by P30 (Figure 2C). The mRNA expression levels of three predominant isoforms of collagen (*COL1A1*, *COL3A1*, and *COL4A1*) were also not downregulated until 6mo in pig left ventricles (Figure 2D). Interestingly, the immature collagen isoform *COL3A1*^24^ is still highly expressed at 2mo in pigs. Conversely, in mice, fibrillar collagen isoforms are significantly downregulated by P30, with highest postnatal *Col3a1* expression at P0-P7 (Figure S3B). Maturation of collagen triple helical fibers was assessed by staining with fluorescent Collagen Hybridizing Peptide (CHP), which specifically binds to denatured and remodeling collagen (Figure 2E-F). In pigs, CHP staining revealed two periods of significant collagen remodeling at P7 and 6mo, compared to P0. In mice, as expected, there is increased trend for collagen remodeling by P7 (Figure S3C). Also, by visual comparison of pigs and mice, CHP morphological expression in pigs at 2mo-6mo appeared similar to fibrillar collagen deposition at P15-P30 in mice (Figure S3A, S3C). This suggests postnatal collagen fiber deposition is robust by P7 in pigs, but with maturational remodeling continuing up to 2mo-6mo, in contrast to the P0-P15 period of ECM maturation in mice.

### 3.3 Cardiac gap junctional maturation and downregulation of fetal contractile protein isoforms occurs after P30 in pigs

In rodents, CM sarcomeres undergo gap junctional maturation in the first two weeks after birth,^25, 26^ identified by colocalization of Connexin-43 (Cx43) to intercalated discs as a measure of CM coupling. In postnatal pigs, immature ‘punctate’ staining of Cx43 along the length of CMs could be seen at P0-P7 (Figure 3A). While terminal localization of Cx43 in CMs was initially detected at P30, complete junctional maturation was only apparent by 2mo-6mo. This is in contrast to mice, in which Cx43 becomes localized at terminal gap junctions by 2 weeks after birth (Figure S4A). With maturation of the CM contractile apparatus, fetal contractile protein isoforms are also downregulated in the first week after birth in mice.^9, 26^ Therefore, the switch from fetal to mature contractile protein isoform switching was measured in pig postnatal ventricular mRNA. Note that in pigs, like humans, MYH7 is the predominant myosin heavy chain isoform in the slowly-contracting mature ventricle, whereas MYH6 predominates in adult mice due to their higher rates of ventricular contraction.^9^ Fetal contractile protein isoforms *TNNI1* and *MYH6* were still detectable at P30 and 2mo respectively in postnatal pigs, while significant upregulation of mature contractile protein isoforms *TNNI3* and *MYH7* did not occur until 6mo (Figure 3B-C). In contrast, fetal contractile protein genes (*Tnni1* and *Myh7*) are completely repressed by P15 in mice, with concurrent upregulation of mature sarcomeric protein genes (*Tnni3* and *Myh6*) (Figure S4B-C). Together, this indicates a P30-6mo window for terminal CM sarcomeric maturation in pig hearts, compared to the P15-P30 time period in mice.

**Figure 3.**
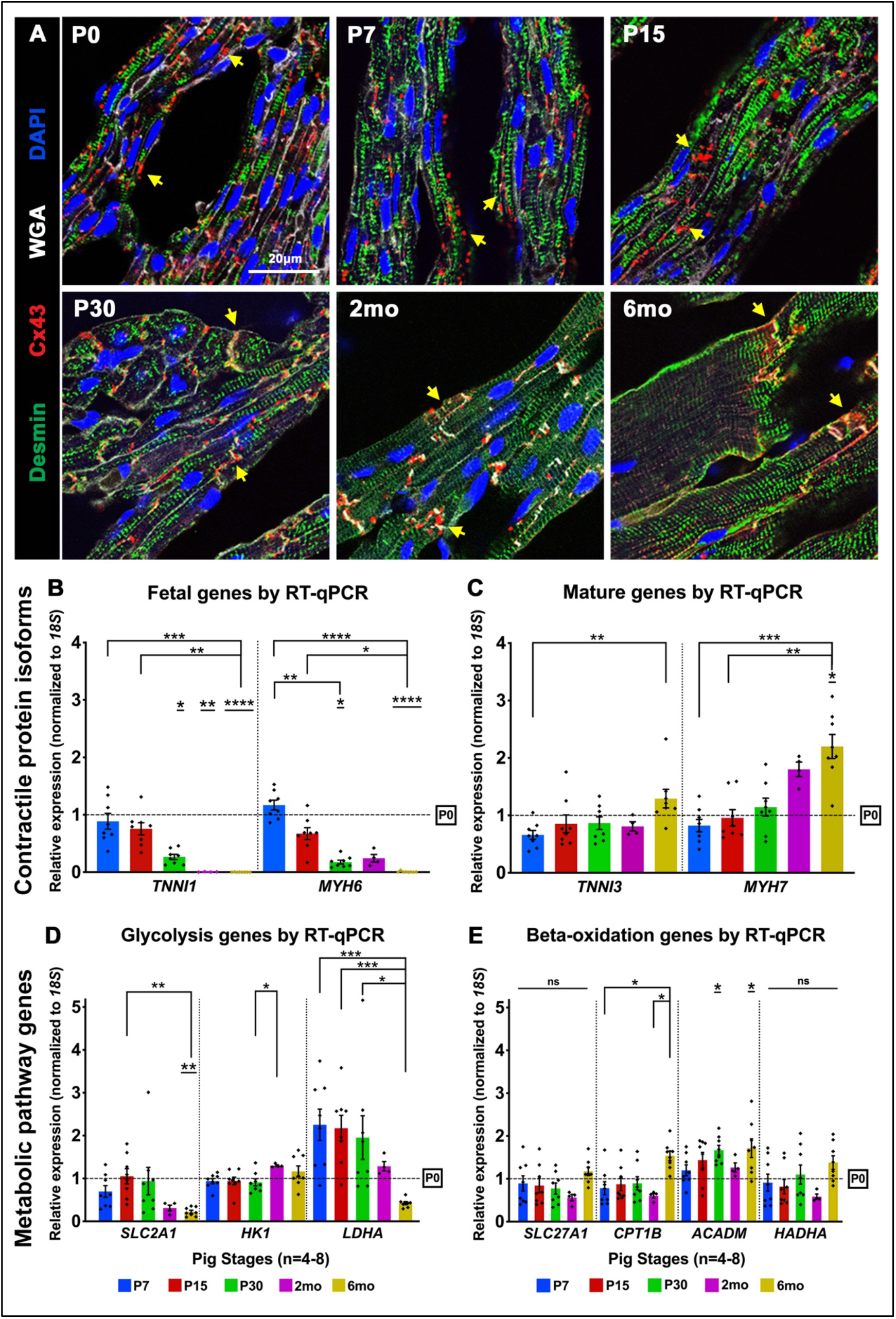
Sarcomeric maturation occurs beyond P15 while no clear transition to betaoxidation metabolism is evident in postnatal pigs. **(A)** Representative images of Connexin-43 (Cx43) staining assessing gap junctional maturation. Yellow arrows show change in Cx43 expression from lateral ‘punctate’ staining at P0-P15 to co-localization at cardiomyocyte termini at P30-6mo. **(B, C)** Expression of cardiac contractile protein fetal and mature isoforms at P7-6mo in pig left ventricles, with fold change relative to P0. **(D, E)** RT-qPCR analysis showing fold change relative to P0 of fetal glycolysis and mature beta-oxidation metabolism genes. Data are mean ± SEM, with *p<0.05, **p<0.01, ***p<0.001, ****p<0.0001 determined by Dunn’s Kruskal-Wallis Multiple Comparisons Tests, in n=4-8 pigs per stage. Asterisk(s) with underline indicate significance compared to P0.

### 3.4 Postnatal pig hearts express glycolytic and beta-oxidation genes

Soon after birth, the neonatal mouse heart transitions from a fetal anaerobic glycolysis metabolic program to mature beta-oxidation metabolism.^8^ Exposure to the high oxygen environment at birth has been implicated in postnatal CM cell cycle arrest and loss of regenerative potential in mice.^27, 28^ Hence, postnatal transitions in metabolic gene expression were examined in pig left ventricular mRNA from P0-6mo. While *SLC2A1* and *LDHA*, two important genes for glycolysis, were significantly downregulated by 2mo-6mo in pig hearts, another key glycolytic gene, *HK1*, remained virtually unaltered from birth to 6mo (Figure 3D). Similarly, when relative mRNA expression of beta-oxidation genes was measured, there was minimal change in expression of three key genes, *SLC27A1*, *ACADM*, and *HADHA* from P0 to 6mo, but *CPT1B* was significantly upregulated by 6mo (Figure 3E). In contrast, in mouse ventricular mRNA, glycolysis genes (*Slc2a1*, *Hk1*, *Ldha*) are downregulated while beta-oxidation genes (*Slc27a1*, *Cpt1b*, *Acadm*, *Hadha*) are all upregulated, in the first week after birth (Figure S4D-E). Thus, in comparison, pig hearts do not have coordinated upregulation of beta-oxidation gene expression after birth. But it is possible that cardiac transition to beta-oxidation metabolism occurs *in utero* in swine, as seen in other large mammal models, such as sheep.^29^

### 3.5 Pig cardiomyocytes are predominantly mononucleated until P15, with extensive multinucleation between P30 and 6mo

In mice, CM binucleation begins in the first week after birth, and nearly all CMs are binucleated by P15, which is maintained in the adult murine heart.^6^ While existence of multinucleated CMs have been reported in young-adult pigs,^15^ the developmental timing and pattern of CM transition to a multinucleated state in postnatal pigs, particularly in the first few months after birth, is not well-defined. We thus assessed CM nucleation in dissociated CMs isolated from pig left ventricles at P0 to 6mo. Individual CMs dissociated from formalin-fixed left ventricular tissue were stained with α-actinin and DAPI for quantification of nuclear number per striated CM (Figure 4A-B). At birth, pig CMs are largely mononucleated (~90%), with a small population having two nuclei per CM, indicating perinatal onset of binucleation in pig hearts. Interestingly, from P0-P15, there remains a significant population of mononucleated CMs, with nearly 50% CM mononucleation evident at P15. Also, loss of CM mononucleation does not occur until 2mo-6mo in pig hearts, with a small population of mononucleated CMs (~10%) seen at P30-2mo. Tetranucleated CMs (4 nuclei per cell) were first observed at P15, representing approximately 30% of all CMs by P30 and 40% by 6mo. Cardiac multinucleation continues progressively in postnatal pigs, with CMs having 8-16 nuclei per cell prevalent by 6mo (~50%). In tissue sections of older pig hearts at 2mo and 6mo, multinucleated CMs could also be seen when stained with WGA to define CM boundaries (Figure 4C). Pig hearts thus show extensive CM multinucleation at P30-6mo, with about 50% mononucleated CMs still present at P15 and persisting at ~10% until 2mo.

**Figure 4.**
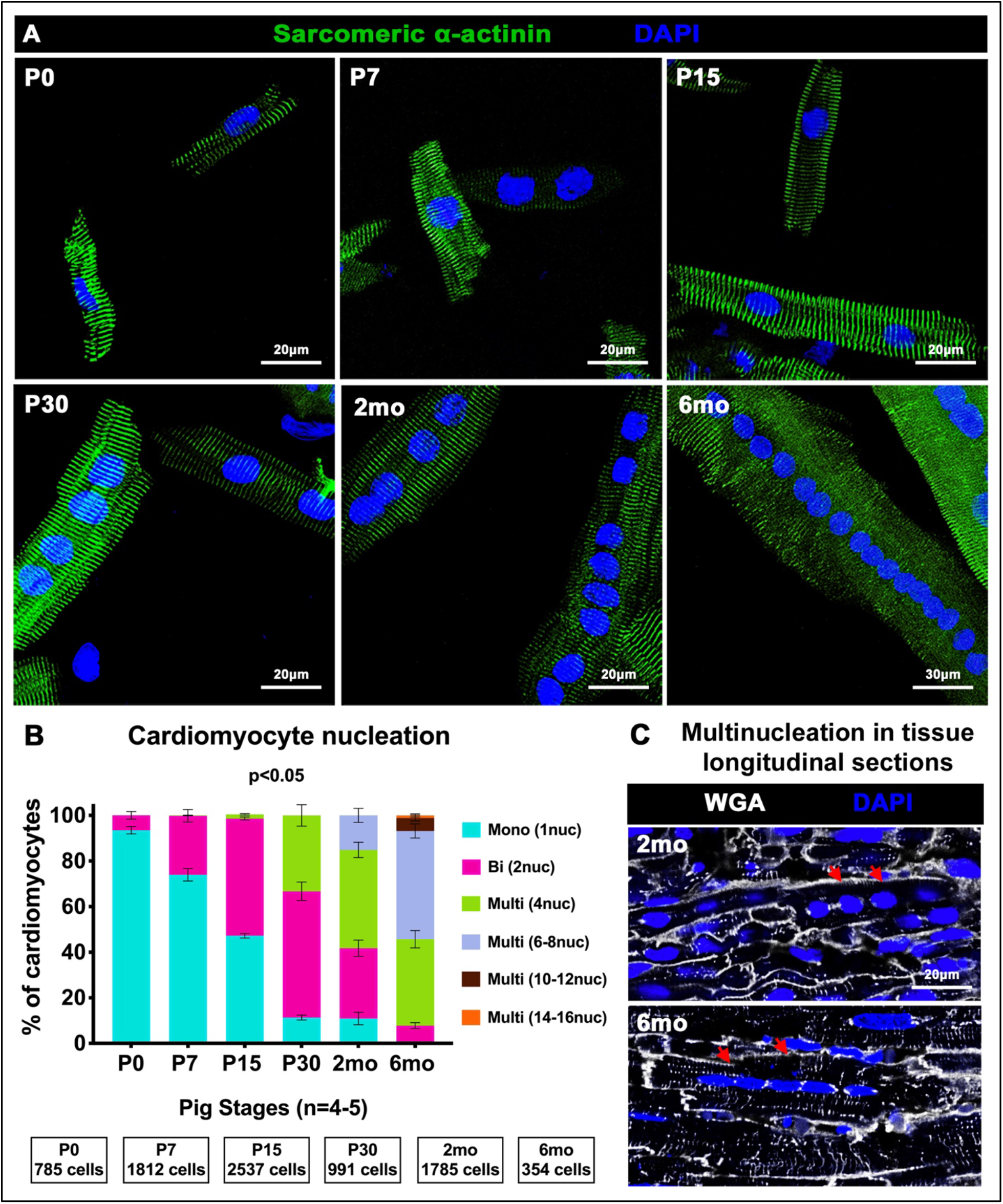
Progressive multinucleation occurs from P0 to 6mo with mononucleation up to 2mo in postnatal pig cardiomyocytes. **(A)** Representative images of pig dissociated CMs from P0 to 6mo showing increased nuclear number indicated by DAPI (blue) in CMs identified by sarcomeric α-actinin (green). **(B)** Number of nuclei per individual CM was determined and percentage of total CMs exhibiting mono-, bi-, and multi-nucleation is indicated for each stage. **(C)** Representative images of multinucleated CMs (red arrows) visualized by WGA staining in longitudinal sections of pig left ventricular tissue. Data are mean ± SEM, with p<0.05 determined by Dunn’s Kruskal-Wallis Multiple Comparisons Test in n=4-5 pigs per stage.

### 3.6 Longitudinal growth with multinucleation precedes age-dependent diametric growth in postnatal pig cardiomyocytes

A characteristic event in the transition of the proliferative neonatal heart to a terminally-differentiated adult heart in rodents is the switch to hypertrophic CM growth.^7^ To evaluate when CM hypertrophic growth is initiated in pig hearts, CM area was measured in left ventricular paraffin sections and in dissociated heart cell preparations. Cross-sectional area (CSA), measured in pig left ventricular tissue stained with WGA and DAPI, revealed no significant change in CM circumference between P0 and P30 in pigs (Figure 5A-B). By 2mo, CSA showed a trend towards increase, which was significant by 6mo. When total cell surface area (TSA) was measured in 2D images of dissociated CMs stained with α-actinin, there was no significant increase from P0 to P15 in CM surface area (Figure 5C-D). Interestingly, TSA showed a trending increase relative to multinucleation (4 or more nuclei per cell) of individual CMs at P15, with significant increase at P30 and beyond. In addition, separate measurements of length and width in dissociated CMs showed longitudinal CM growth in bi- and multi-nucleated CMs at all stages, with the extent of increase in CM length proportional to number of nuclei per cell. However, increased CM width was only evident by 2mo-6mo (Figure 5E-F). Further, we confirmed technical consistency of our CM cytoplasmic area (CSA) measurements in paraffin sections by contrasting against CM nuclear surface area measured in the same cross-sectional region (Figure 5G). Average area per nucleus does not change across stages while average cytoplasmic area per CM increases with age, confirming our observations of age-related width expansion of CMs in pigs. Together, these data demonstrate a differential mode of CM growth in pigs compared to mice, with longitudinal CM growth occurring relative to multinucleation beyond P15 in postnatal pigs, and diametric CM growth having significantly increased cross-sectional area and width measured at 2mo-6mo.

**Figure 5.**
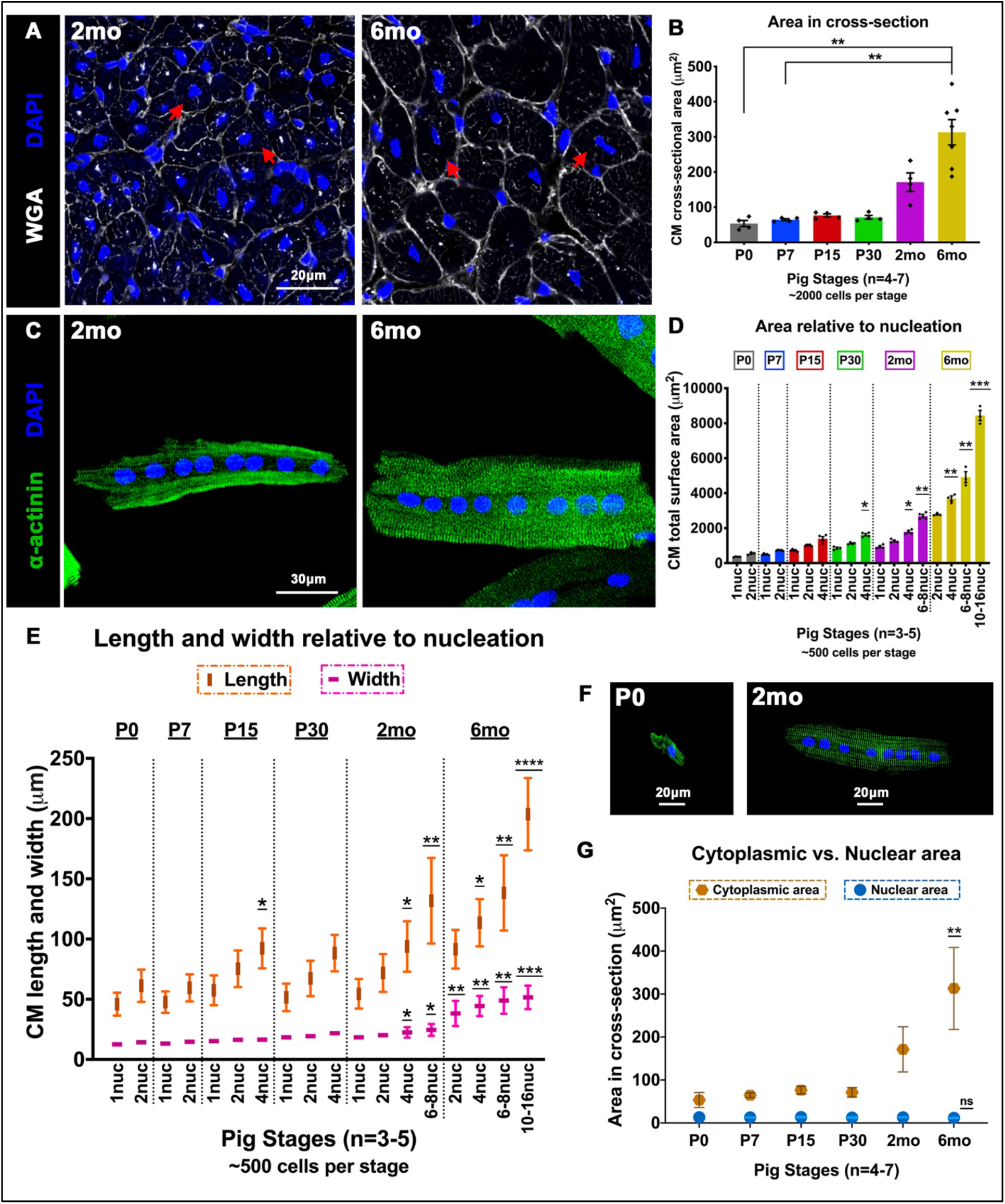
Cardiomyocyte length increases relative to nucleation from P0-6mo while width does not increase until 2mo-6mo in postnatal pig hearts. **(A)** Representative images of pig heart cross-sections stained with WGA and DAPI at 2mo and 6mo. Red arrows indicate increase in CM cross-sectional cell size. **(B)** CM cross-sectional area (μm^2^) in paraffin sections was quantified by manual cell tracing based on WGA staining. **(C)** Representative images of dissociated pig CMs showing increased width between octa-nucleated cells at 2mo and 6mo **(D)** Total surface area (μm^2^) was measured by manual cell tracing of dissociated CMs based on α-actinin staining. **(E)** Length and width were assessed in dissociated pig CMs by manual measurements. **(F)** Representative images of change in CM length relative to nucleation at P0 and 2mo. **(G)** Nuclear area based on DAPI staining was measured and compared against cytoplasmic area measured by cell tracing of WGA staining in paraffin cross-sections. Data are mean ± SEM in B and D, and mean ± SD in E, with *p<0.05, **p<0.01, ***p<0.001, ****p<0.0001 determined by Dunn’s Kruskal-Wallis Multiple Comparisons Tests. Asterisk(s) indicate significance compared to mononucleated P0 CM (P0-1nuc) in D and E. In G, data are mean ± SD, with **p<0.01 and ns=not significant determined compared to P0 by Brown-Forsythe and Welch ANOVA tests with Games-Howell corrections for Multiple Comparisons (n=3-7 pigs per stage).

### 3.7 Pig cardiomyocyte nuclei are primarily diploid in the 6 months after birth

Postnatal CM polyploidization has been proposed as a mechanism for loss of proliferative capacity in the adult mammalian heart.^12, 14^ In mice, CM polyploidization occurs by binucleation, with individual CM nuclei remaining predominantly diploid (2c).^12^ Meanwhile, humans exhibit mononucleated CMs with polyploid (>2c) nuclear DNA content.^30^ In pigs, overall CM polyploidization occurs by progressive multinucleation from P0-6mo (Figure 4), however, ploidy per nucleus is unknown. Hence, to assess the DNA content of individual nuclei in pig CMs, Hoechst staining was quantified. In dissociated cell preparations, CM nuclear Hoechst intensity was normalized to haploid (1c) sperm nuclei per image (Figure 6A-B). The relative intensity of CM nuclei to sperm nuclei did not significantly change across all stages (Figure 6B). Assuming the normalized value for P0 mononucleated CM nuclei indicates 2c DNA content, this suggests that individual CM nuclei remain diploid from P0 to 6mo in pigs. Interestingly, there is also no significant change in nuclear DNA content of mononucleated CMs from P0 to 2mo, suggesting the ploidy of mononucleated CMs does not change after birth in pigs. Following this, Hoechst intensity was also measured in left ventricular cryosections co-stained with Desmin (CM) and Vimentin (non-CM) to assess nuclear DNA content, using Vimentin-positive cell nuclei as 2c control for normalization per image (Figure 6C-D). Thresholds for 2c and >2c cut-offs were determined as described in supplementary methods. There was no significant change in normalized Hoechst intensities from P0 to 6mo, indicating individual pig CM nuclei are mostly diploid in the first six postnatal months. When percentages were estimated, approximately 90% of CM nuclei were within the normalized threshold for diploid DNA content (Figure 6E). These percentages are similar to a previously reported study, which found 75%-85% diploid CM nuclei in juvenile and adult pig hearts.^31^ Further, measuring nuclear Hoechst intensity in multinucleated CMs within longitudinal sections of cardiac tissues also revealed primarily diploid nuclei, as well as consistent DNA content in individual nuclei of the same multinucleated CM (Figure 6F-G). This supports our observation in dissociated pig CMs (Figure 6A-B) that there is no change in DNA content per nucleus in multinucleated CMs after birth. Together, these data show the predominance of diploid (2c) DNA content per nucleus in mono-, bi- and multi-nucleated CMs from P0-6mo in pigs.

**Figure 6.**
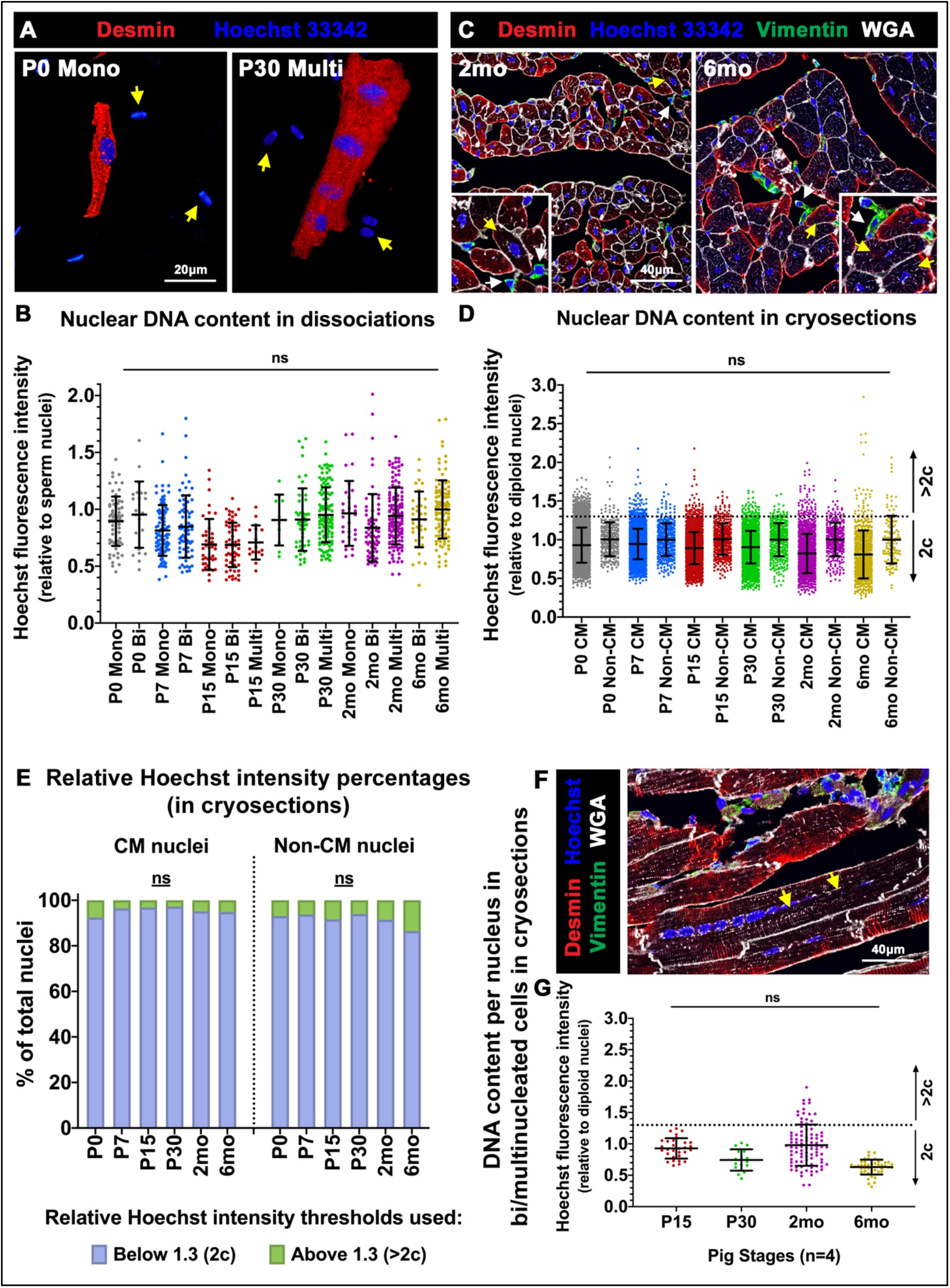
Cardiomyocyte nuclei remain predominantly diploid in postnatal pig hearts (P0-6mo). **(A)** Representative images of dissociated CMs stained with Hoechst 33342 for nuclear intensity assessment, with sperm nuclei (yellow arrows) utilized for normalization. **(B)** Hoechst fluorescence intensity per nucleus is represented relative to sperm nuclear intensities per image (>100 CMs counted per stage). **(C)** Representative images of ventricular tissue cryosections stained with Hoechst 33342 for CM nuclear intensity assessment (yellow arrows), with Vimentin-positive cell nuclei (white arrows) utilized as diploid (2c) control. **(D)** Hoechst fluorescence intensity per nucleus is represented relative to Vimentin-positive cell nuclear intensities per image. Thresholds for 2c and >2c were determined based on standard deviation of non-CM (2c) nuclei (>100 CMs counted per stage). **(E)** Percentage of CM and non-CM nuclei with relative Hoechst intensities at 2c or >2c, as determined in ventricular tissue cryosections. **(F)** Representative image of a multinucleated CM (yellow arrows) in longitudinal tissue section stained with Desmin, Hoechst, and WGA for nuclear intensity measurements. **(G)** Hoechst fluorescence intensity per nucleus in multinucleated CMs was measured at P15-6mo in cryosections. Data are mean ± SD, with ns=not significant determined compared to P0 by Brown-Forsythe and Welch ANOVA tests, in n=4 pigs per stage.

### 3.8 Cardiomyocytes are mitotically active up to 2mo in pig left ventricles

CM cell cycle arrest and loss of mitotic activity in mouse hearts occurs in the first week after birth.^4–6^ To determine timing of cardiac cell cycle arrest in postnatal pigs, mitotic index and cell cycle gene expression were assessed. Staining of pig heart sections with Phosphohistone-H3 (pHH3), a marker of mitosis, revealed 0.1% of CM nuclei are pHH3-positive up to 2mo in pigs, and there is no significant decrease in pHH3 levels until 6mo compared to neonatal stages (Figure 7A-D). Thus, while CM mitotic activity declines precipitously within a week after birth in mice,^5, 6^ CM mitotic activity in pigs does not decline until after 2mo in the postnatal period. The highest pHH3 levels (~0.3% of CM nuclei) were measured at P7 and P15 in pigs, likely in-part due to onset of bi- and multi-nucleation at these stages. Interestingly, when longitudinal sections of 2mo pig hearts were examined, simultaneous activation of pHH3 in adjacent nuclei of multinucleated CMs could be detected, indicating karyokinesis in the absence of cytokinesis (Figure 7C). Thus, mitotically-active CMs are present up to 2mo in pig hearts, with karyokinesis in the absence of cytokinesis apparent in multinucleated CMs in older pigs.

Cell cycle gene expression was then evaluated by assessment of cell cycle promoting and inhibitory genes in pig left ventricular mRNA (Figure 7E-F). In agreement with pHH3 staining showing mitotic activity up to 2mo (Figure 6A-D), cardiac cell cycle gene expression is evident up to 2mo in pigs, and downregulation of cell cycle promoting genes does not occur until 6mo (Figure 6F). While G1 (Growth) phase genes remain largely unchanged after birth in pig left ventricles, S- (DNA synthesis) and M- (Mitosis) phase genes exhibit no significant decrease in expression from P0 to P30. Also, interestingly, among M-phase genes, *PRC1* (a cytokinetic regulator) is not significantly decreased until after 2mo in postnatal pig hearts. Cell cycle inhibitor genes for S- and M-phases are also not significantly upregulated until 6mo in pig hearts. This is in contrast to mice, where cell cycle promoting *Cyclins* and *CDKs* are significantly downregulated beyond P7 (Figure S5), compared to P0 levels. Further, developmental and proliferative cardiac signaling pathways such as AKT and β-catenin signaling, reduced soon after birth in mice,^32^ are not downregulated until 6mo in pig left ventricular protein as determined by Western blotting (Figure S6). Thus, these results indicate the presence of actively-mitotic CMs until 2mo in pig hearts, with cardiac cell cycle arrest and repression of developmental growth pathways not occurring until 6mo.

**Figure 7.**
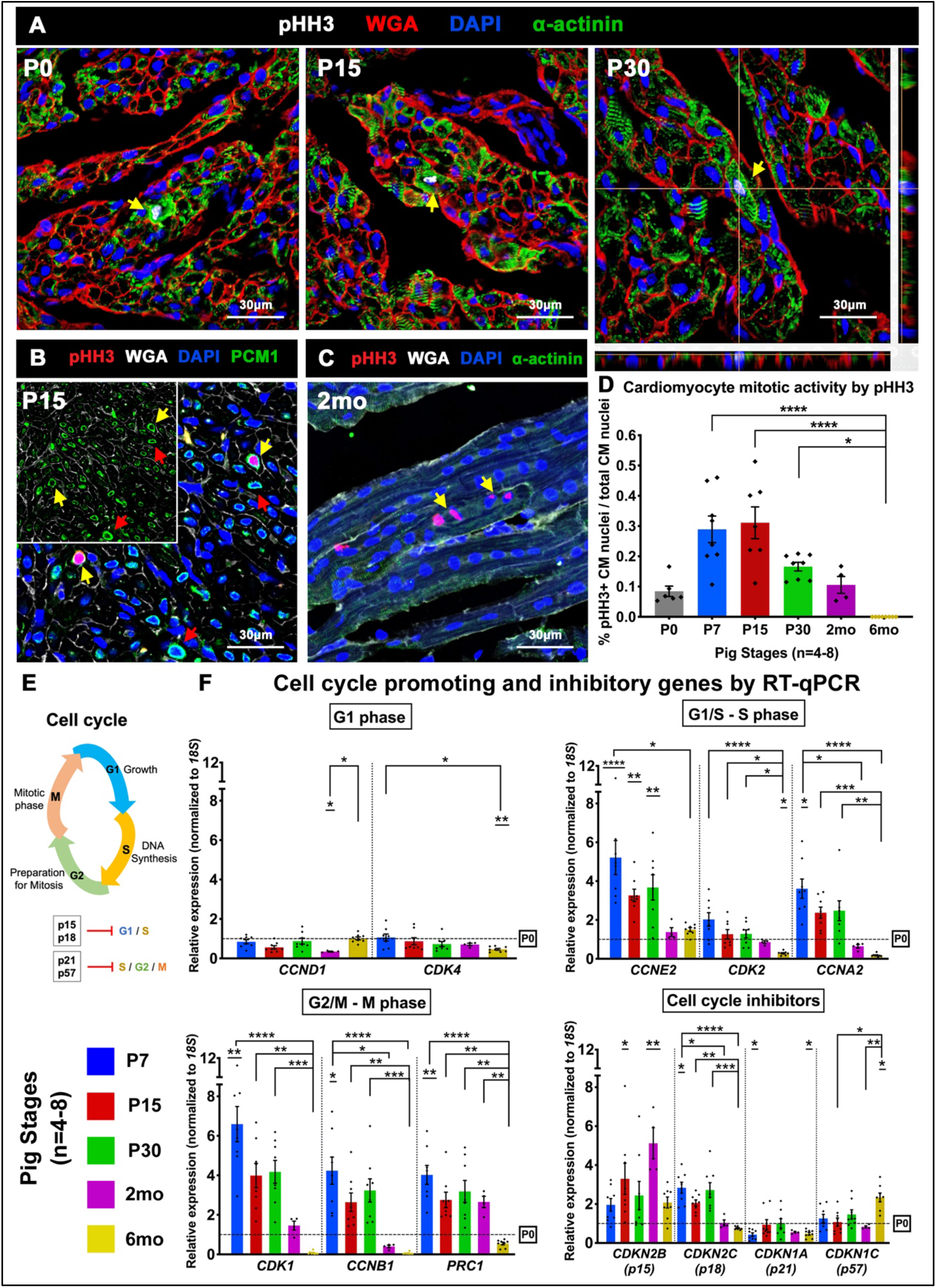
CM mitotic activity and cell cycle gene expression are detected up to 2mo in postnatal pig hearts. **(A, B)** Representative images of Phosphohistone-H3 (pHH3) staining for mitotic activity (yellow arrows) in pig heart tissue sections. CMs were identified by sarcomeric α-actinin (A, green stain) and perinuclear PCM1 (B, red arrows). **(C)** Longitudinal tissue sections showed presence of multinucleated CMs with simultaneous pHH3 activation in adjacent nuclei (yellow arrows). **(D)** Ratio of pHH3-positive CM nuclei to total CM nuclei was calculated in cardiac cross-sections. **(E)** Schematic of cell cycle phases. **(F)** RT-qPCR analysis for cell cycle promoting and inhibitory genes was assessed, with fold change relative to P0. Data are mean ± SEM, with *p<0.05, **p<0.01, ***p<0.001, ****p<0.0001 determined by Dunn’s Kruskal-Wallis Multiple Comparisons Tests in n=4-8 pigs per stage. Asterisk(s) with underline indicate significance compared to P0.

## 4. Discussion

In this study, we show that the major events of postnatal cardiac maturation are delayed in pigs relative to mice, and that pig CMs display growth dynamics not seen in other mammals (Figure 8). Postnatal cardiac maturation, including collagen deposition, sarcomeric maturation, loss of mononucleated-diploid CMs, and cardiac cell cycle arrest, occurs over a 2mo-6mo period in pig hearts, compared to the 1-2 week period of terminal cardiac maturation after birth in mice.^20^ Further, pigs exhibit possibly-unique postnatal CM growth characteristics among mammals, as evidenced by extensive CM multinucleation beyond P30, with up to 16 centrally-located nuclei in a single CM seen at 6mo. In addition, pig CMs initially grow longitudinally during multinucleation, and conventional hypertrophic growth as indicated by increased cross-sectional area is only induced several months after birth. However, despite such fundamental differences in the timing and pattern of heart maturation, cardiac regenerative potential only extends up to the first week after birth in both mice and pigs.^16–18^ This discordant timing of postnatal myocardial maturational events in pigs relative to the time of loss of heart regenerative capacity may thus provide novel insights into the underlying mechanisms of mammalian cardiac regeneration.

**Figure 8.**
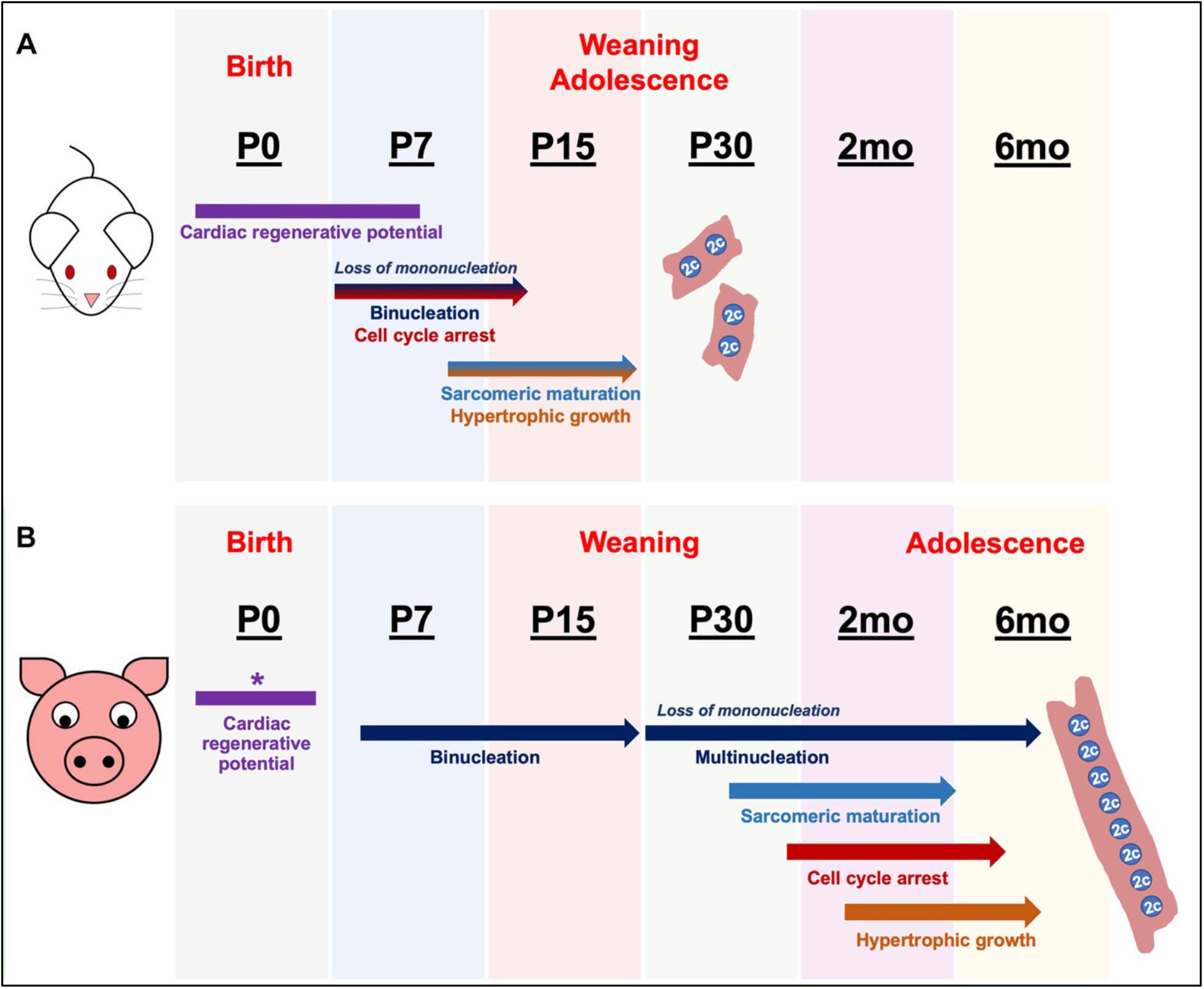
Delayed cardiac terminal maturation despite similar neonatal periods of heart regenerative potential in pigs compared to mice. **(A)** Schematic of postnatal developmental events in murine hearts, showing loss of cardiac regenerative capacity at P7 coincident with CM cell cycle arrest, binucleation, and maturation. **(B)** Schematic summarizing postnatal pig cardiac developmental events, showing discordant timing of cardiac maturational milestones in pigs compared to mice. *Pig cardiac regenerative window is as described in previous reports.^17, 18^

### 4.1 Postnatal cardiomyocyte growth dynamics are distinctive in swine compared to other mammals

In rodents, CMs are primarily binucleated by P15, with each nucleus generally being diploid.^7^ In humans, while the prevalence of binucleated CMs is debated, it is commonly accepted that adult CMs are mainly mononucleated with polyploid DNA content per nucleus with increasing age.^13, 33, 34^ When other mammals such as dogs, cats, and sheep are considered, only bi-, tri- or tetra-nucleation (2-4 nuclei per CM) is evident.^11^ Even outside of mammals, across 41 diverse vertebrate species, only up to tetranucleated CMs was reported.^14^ In pigs, however, while onset of binucleation occurs around birth, extensive multinucleation of 4, 8, and up to 16 nuclei per CM is observed by 6mo, with individual nuclei remaining largely diploid. Up to 32 nuclei per CM has previously been recorded in older swine.^15^ The role of such extensive multinucleation and its implications on cardiac function in pigs are unknown, but from our data, multinucleation contributes to longitudinal elongation of CMs in the rapidly-growing hearts of young swine.

The switch to a hypertrophic mode of CM growth is an important landmark for terminal maturation of the mammalian heart.^7, 10, 13, 35^ In pigs, this transition occurs in a distinctive pattern, where CM length relative to nucleation is increased prior to the increase in CM width relative to age. This is a possibly-unique mechanism of CM hypertrophy among mammals. The only other study reporting a similar mode of heart muscle growth among vertebrates is in chicken, where a small percentage of multinucleated CMs (>5 nuclei per CM) was reported during longitudinal CM growth, both preceding the increase in CM width.^36^ It is possible that farm animals like chickens and pigs, selectively bred for meat production, utilize growth by CM length and multinucleation to achieve the high cardiac demands of rapid postnatal body mass increase. Interestingly, increased muscle fiber length through multinucleation is widely considered a skeletal muscle-like growth mechanism,^37^ and in skeletal muscle cells, multinucleation occurs through syncytial fusion, with nuclei localized towards the surface of individual myofibers. However, individual pig CM nuclei are arranged linearly along the central axis of the cell, with multinucleation occurring via karyokinesis. This presents intriguing questions about the structural and physiological differences of pig CMs compared to other mammalian CMs, which will need to be explored to understand the role played by multinucleation in pig CM functionality. Further, conventional measurements of CM hypertrophy only assess cross-sectional area (i.e. width).^7^ Thus, the prolonged period of increased CM longitudinal growth alongside later induction of diametric growth in swine may be useful to elucidate novel mechanisms of non-proliferative heart growth.

### 4.2 Cardiac terminal maturational events extend to 2-6 postnatal months in pigs, beyond the P3 regenerative period

In pigs, cardiac terminal maturation as defined by cell cycle exit, sarcomeric maturation, loss of mononucleation, and complete transition to hypertrophic growth occurs over a 2 to 6-month postnatal period. However, recent studies show that pigs only have a P3 neonatal cardiac regenerative window, with injury models at P7 and P14 developing extensive scarring.^17, 18^ Results from our lab also show a lack of regenerative capacity in P30 pig hearts subjected to transient ischemia/reperfusion injury, with scar formation and no increase in CM mitotic activity by one month post-surgery (Agnew *et al*., unpublished results). Analyzing the discordant timing of postnatal cardiac maturational events in pigs could therefore have important insights into the mechanisms regulating loss of heart regenerative potential after birth in mammals.

Lack of mononucleated-diploid and cell cycling CMs has been proposed as a mechanism for loss of cardiac regenerative capacity in adult mammals.^5, 12^ The innate capacity in zebrafish and newts for heart regeneration throughout life has been linked to presence of mononucleated-diploid CMs,^38^ while polyploidization of CM nuclei inhibits cardiac regeneration in zebrafish.^12, 39^ Further, mononucleated-diploid CMs were used as a proxy for regenerative potential in assessing the impact of evolutionary ectotherm to endotherm transition on cardiac regenerative capacity.^14^ Rodent hearts undergo cell cycle arrest in the first postnatal week concurrent with loss of heart regenerative capacity, and cell cycling is also negligible in the non-regenerative adult human heart.^16, 30^ However, we observe mononucleated-diploid CMs up to 2mo in postnatal pigs, with over 50% CMs exhibiting mononucleation at P15. We also show that active cardiac cell cycling and robust CM mitotic activity occur up to 2mo in pig hearts. Hence, there is a lack of regenerative capacity in pig hearts after P3,^17, 18^ despite the significant presence of mononucleated-diploid CMs at P15 and active cardiac cell cycling up to 2mo. Also, karyokinesis in the absence of cytokinesis occurs in multinucleation of pig CMs, suggesting that induction of cell cycle gene expression alone, as has been suggested,^40^ will possibly not support regenerative repair of heart muscle after injury in swine. Exploring the mechanisms driving multinucleation without CM proliferation in the pig model could thus provide unique insights into postnatal CM proliferative capacity and cytokinetic arrest in mammals. Our results also support the importance of factors other than CM cell cycling and mononucleation in determining cardiac regenerative capacity.

### 4.3 What dictates postnatal regenerative capacity of the mammalian heart?

Several factors in addition to CM proliferative capacity have been implicated in heart regenerative potential. These include maturation of cardiac fibroblasts, activity of macrophages and other immune cells, ECM composition and rigidity, as well as physiological hypoxia and hormonal regulation.^14, 28, 41, 42^ Investigating the discordance in timing of cardiac maturation and regenerative capacity in pigs could be useful to define relative contributions of multiple cell types and physiological factors in heart regeneration. Here, we show that ECM remodeling is significant by P7 in pigs, contributing to increased rigidity of the matrix, while maturational collagen remodeling continues to 6mo. This increase in collagen after birth could contribute to loss of regenerative capacity in swine, as has been proposed in rodents.^23^ Likewise, perinatal hypoxia and oxidative stress have also been linked to loss of heart regenerative capacity after birth in mice,^27, 28^ suggesting that cardiac regeneration can occur only in the first few days after birth in all mammals. Another physiological determinant of cardiac maturation postulated is evolutionary metabolic transition towards endothermy, where thyroid regulation was linked to increased CM ploidy.^14^ Perturbation of thyroid hormone signaling was also found sufficient to enhance CM proliferation. However, multinucleated CMs and large terrestrial mammals, such as pigs, sheep, and cows, were not highlighted in this study.^14^ Maturation of the innate immune system is also a possible mechanism for loss of cardiac regenerative ability in the weeks after birth, but this has not been studied in depth beyond initial reports in rodents.^42^ Together, there is increasing evidence that CM cell cycle arrest is not the only determinative factor in loss of cardiac regenerative potential in mammals after birth. Further studies in pigs will likely be informative in identifying critical parameters for cardiac regeneration in large mammals.

### 4.4 Is there an optimal mammalian model for preclinical studies of human heart disease?

Juvenile pigs at 2mo-6mo are commonly utilized in preclinical testing of cardiac injury mechanisms and therapies for cardiovascular disease.^3^ As we show here, pig hearts are still actively undergoing maturational remodeling during this postnatal period. Further, our study highlights fundamental variations in pig CM characteristics such as multinucleation and differential mechanisms of hypertrophy and cell cycling compared to rodents and humans. Therefore, it may prove important to consider these differences while designing tests for cardiac regenerative therapeutics using swine as a large mammal model. Ultimately, there may not be a single mammalian model that recapitulates the human heart. However, investigation of such differences in CM nucleation and cell cycling mechanisms in pigs and other mammals could offer unique opportunities to study aspects of CM proliferation and cardiac regeneration, thereby enhancing clinical translatability of therapies for human heart disease.

## Acknowledgements

We thank Dr. Bernhard Kühn (UPMC Children’s Hospital, Pittsburgh) for providing cardiomyocyte dissociation protocols, and Dr. Michaela Patterson (Medical College of Wisconsin, Milwaukee) for critical suggestions in optimizing ploidy assessment in pig cardiomyocytes. We also thank past and current members of the Yutzey Lab for assistance in harvest of pig cardiac tissues as well as valuable discussions.

## Author Contributions

The study was designed by NV, CMA, EJA, and KEY. Execution of experimental procedures was by NV, CMA, EJA, KWR, RSB, and FZ. Data was interpreted by NV, CMA, EJA, and KEY. Figures and manuscript were prepared by NV and KEY. Manuscript was edited and approved for final version by all authors.

