## Supplemental figures, tables and methods for "Cardiomyocyte cell cycling, maturation, and growth by multinucleation in postnatal swine"

**SUPPLEMENTARY INFORMATION**

**Velayutham *et al.,* ‘Cardiac maturation in postnatal swine’**

#### Supplementary Figures

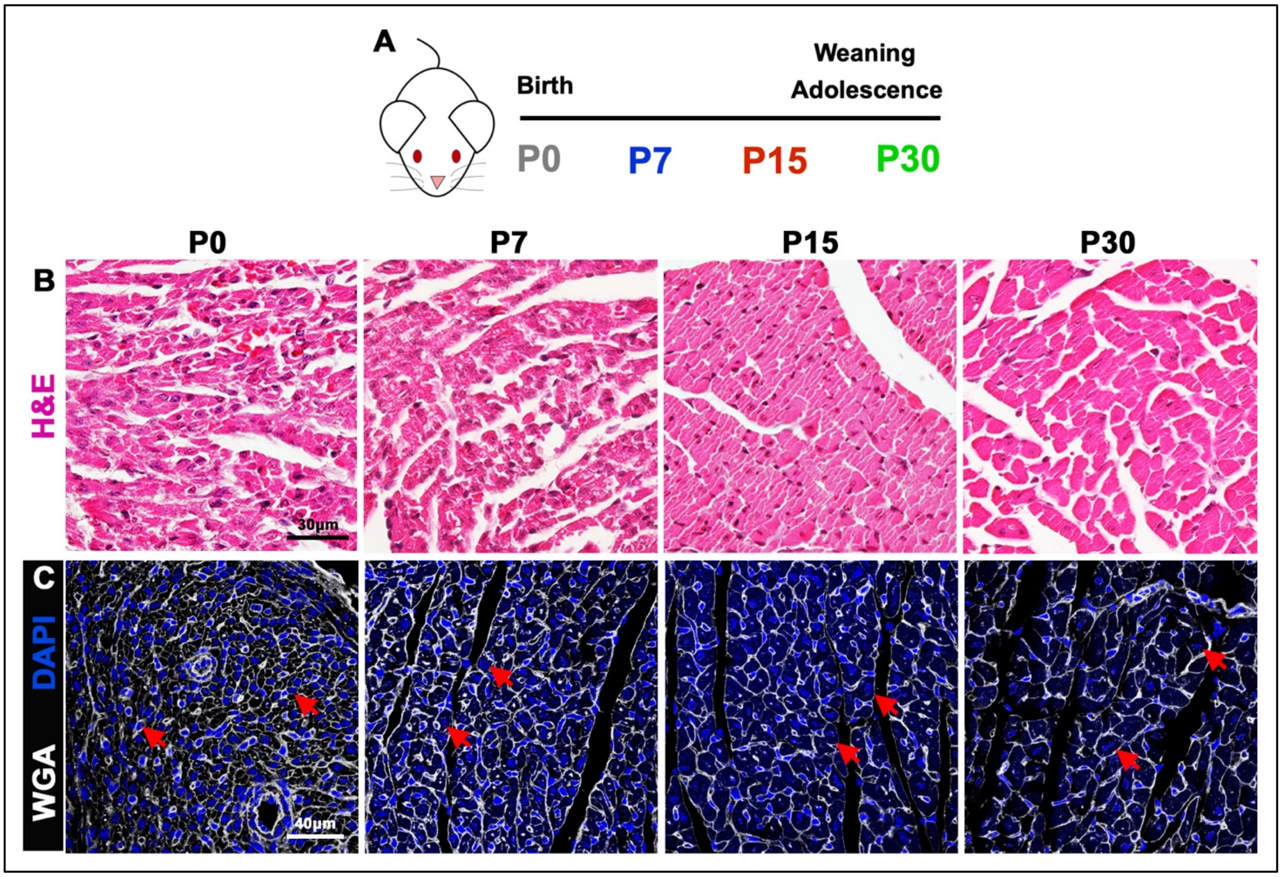


**Figure S1. Postnatal mouse hearts are terminally mature by P30. (A)** Schematic of mouse heart stages utilized for this study. **(B)** Representative images of ventricular morphology visualized by Hematoxylin & Eosin (H&E) and Wheat Germ Agglutinin (WGA) stains (n=3 mice per stage). Red arrows indicate changes in cardiomyocyte cell size in cross-section over the postnatal period.

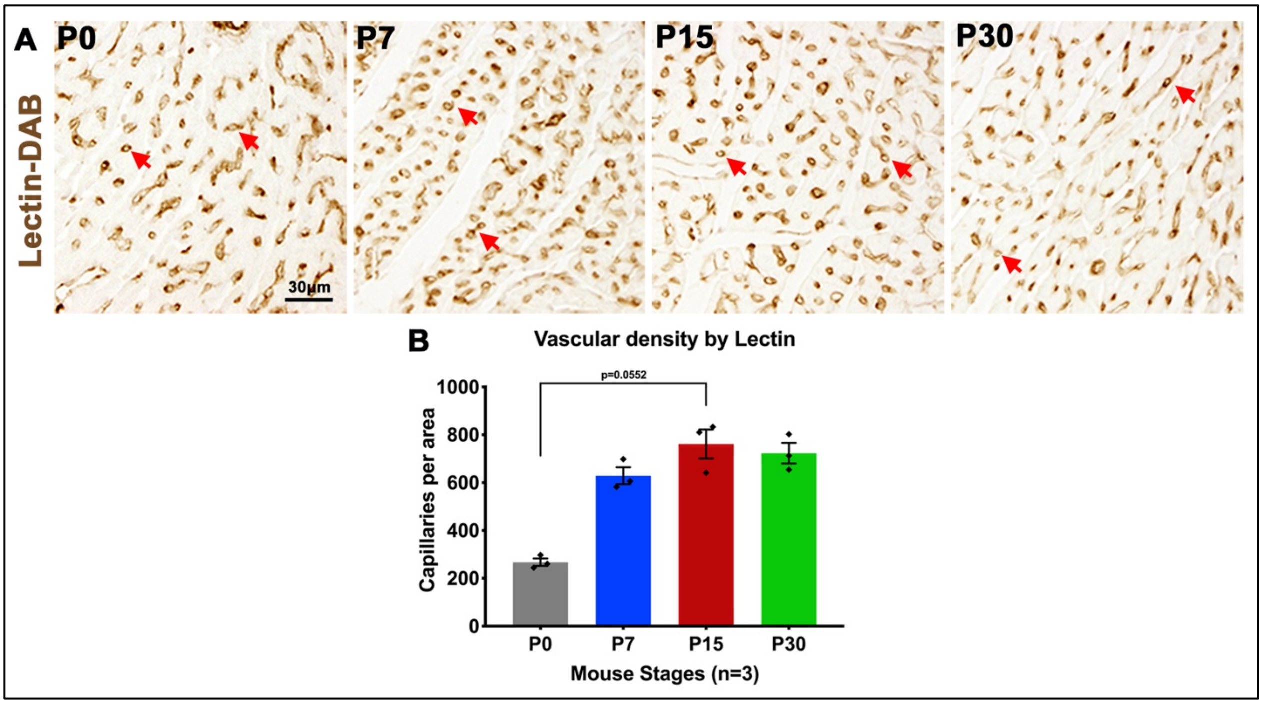


**Figure S2. Increasing trend of vascular density by P7 in postnatal mouse hearts. (A)** Representative images of circular capillaries (red arrows) identified by Lectin-DAB staining in mouse hearts. **(B)** Vascular density was assessed by manual counts for Lectin-DAB positive structures per tissue area ($0.3mm^2$). Data are mean ± SEM, statistical analysis was performed by Dunn's Kruskal-Wallis Multiple Comparisons Test (n=3 per stage).

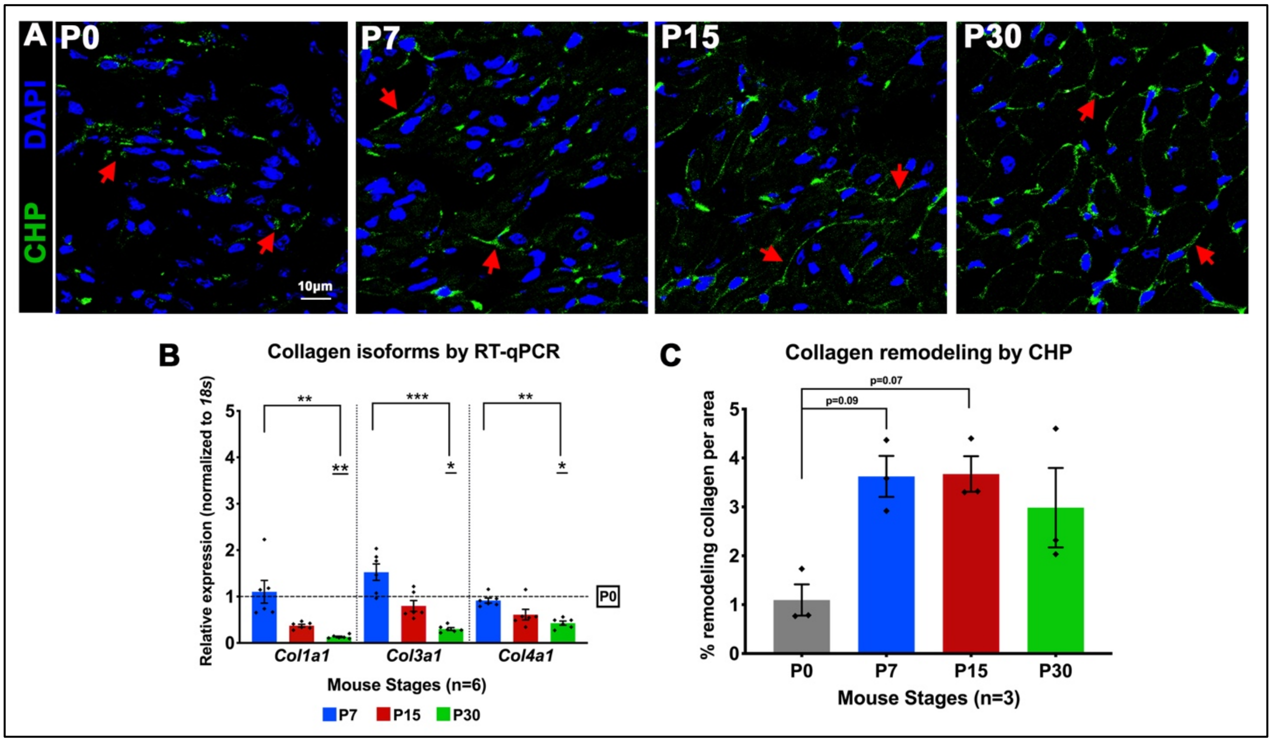


**Figure S3. Increasing trend of collagen remodeling by P7 with downregulation of collagen gene expression by P30 in mouse hearts. (A)** Collagen remodeling was assessed by fluorescent Collagen Hybridizing Peptide (CHP) staining (red arrows). **(B)** RT-qPCR analysis of collagen isoform gene expression, with fold change relative to P0. **(C)** Quantification of collagen remodeling by measuring CHP expression (green staining) per area ($0.05mm^2$). Data are mean ± SEM, with *p<0.05, **p<0.01, ***p<0.001 determined by Dunn's Kruskal-Wallis Multiple Comparisons Tests (n=3-6 per stage). In B, asterisk(s) with underline indicate significance compared to P0.

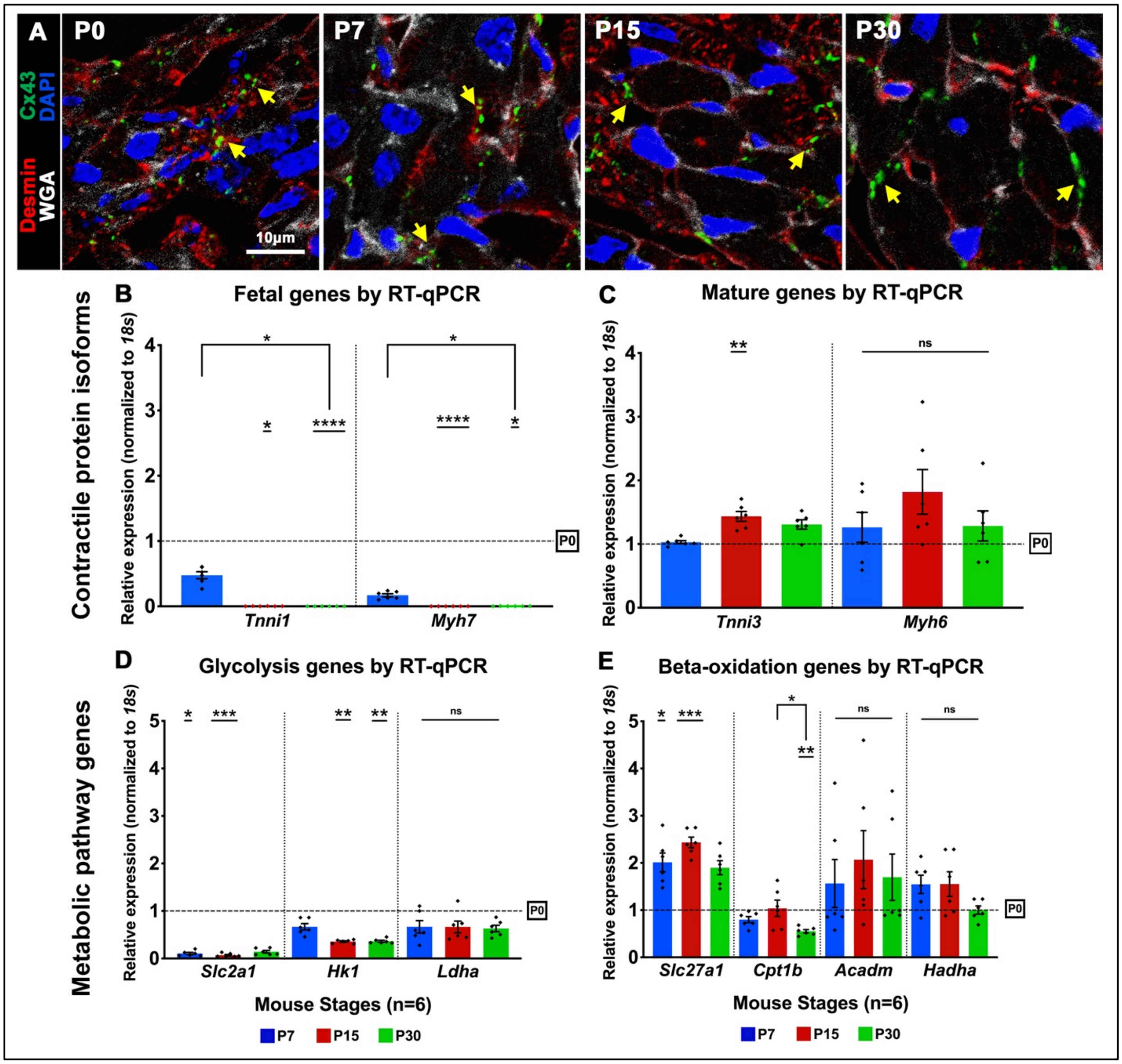


**Figure S4. Sarcomeric maturation and metabolic switching occur by P7-P15 in mouse hearts.** **(A)** Representative images of Connexin-43 (Cx43) staining for gap junctions, showing maturational terminal localization of Cx43 in cardiomyocytes identified by Desmin expression (yellow arrows). **(B, C)** RT-qPCR analysis for cardiac contractile protein isoform switching from fetal (*Tnni1* and *Myh7*) to mature (*Tnni3* and *Myh6*) genes with fold change calculated relative to P0, in mouse ventricular mRNA. **(D, E)** RT-qPCR analysis for metabolic switching from fetal glycolysis to mature beta-oxidation metabolism gene expression with fold change calculated relative to P0, in mouse ventricular mRNA. Data are mean ± SEM, with $^{*}p<0.05$, $^{**}p<0.01$, $^{***}p<0.001$, $^{****}p<0.0001$ determined by Dunn's Kruskal-Wallis Multiple Comparisons Tests (n=3-6 per stage). Asterisk(s) with underline indicate significance compared to P0.

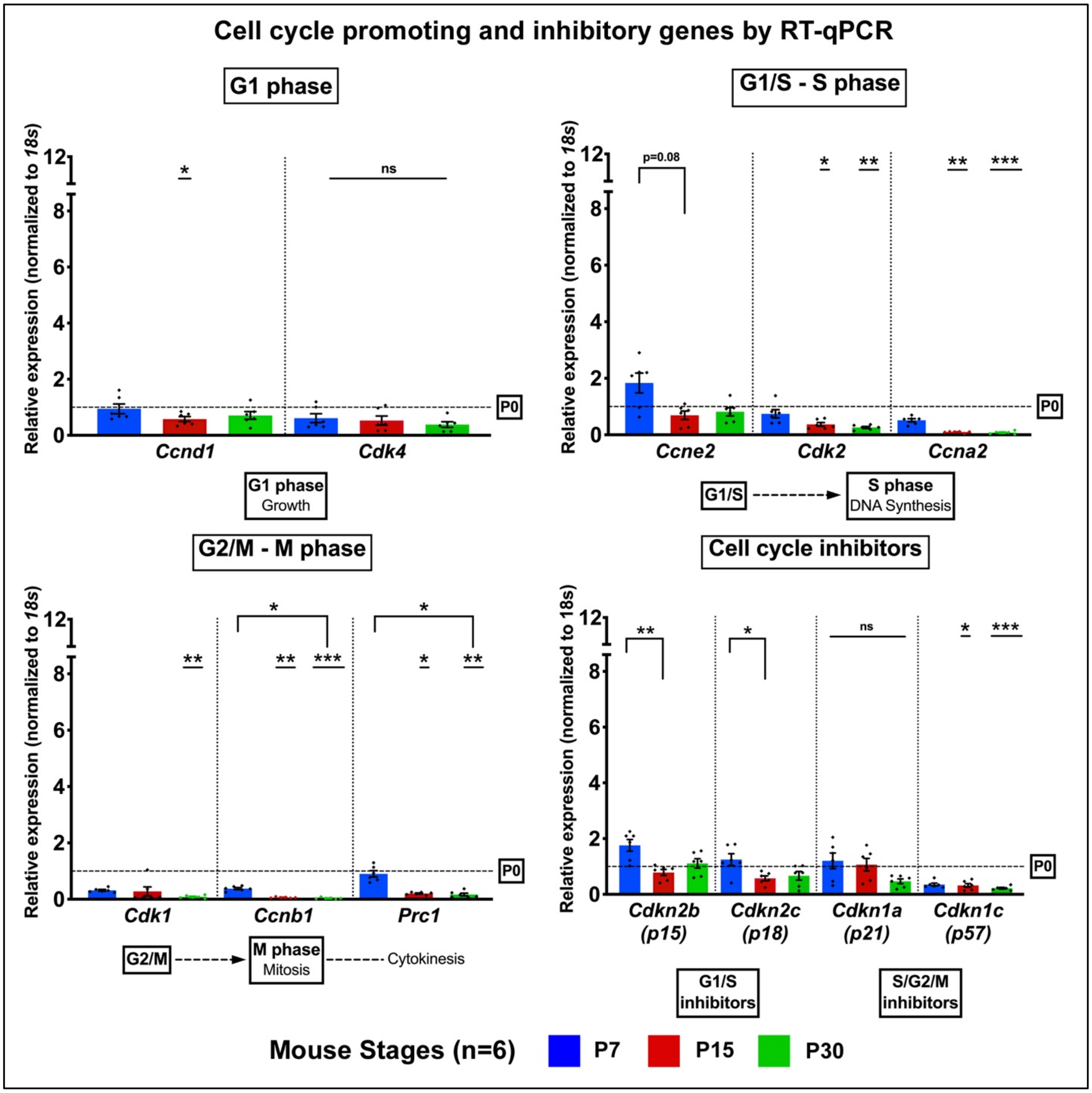


**Figure S5. Repression of cell cycle gene expression occurs by P7-P15 in mouse ventricles.** RT-qPCR analysis for cell cycle promoting and inhibitory gene expression with fold change calculated relative to P0, in mouse ventricular mRNA. Data are mean ± SEM, with *p<0.05, **p<0.01, ***p<0.001 determined by Dunn's Kruskal-Wallis Multiple Comparisons Tests (n=6 per stage). Asterisk(s) with underline indicate significance compared to P0.

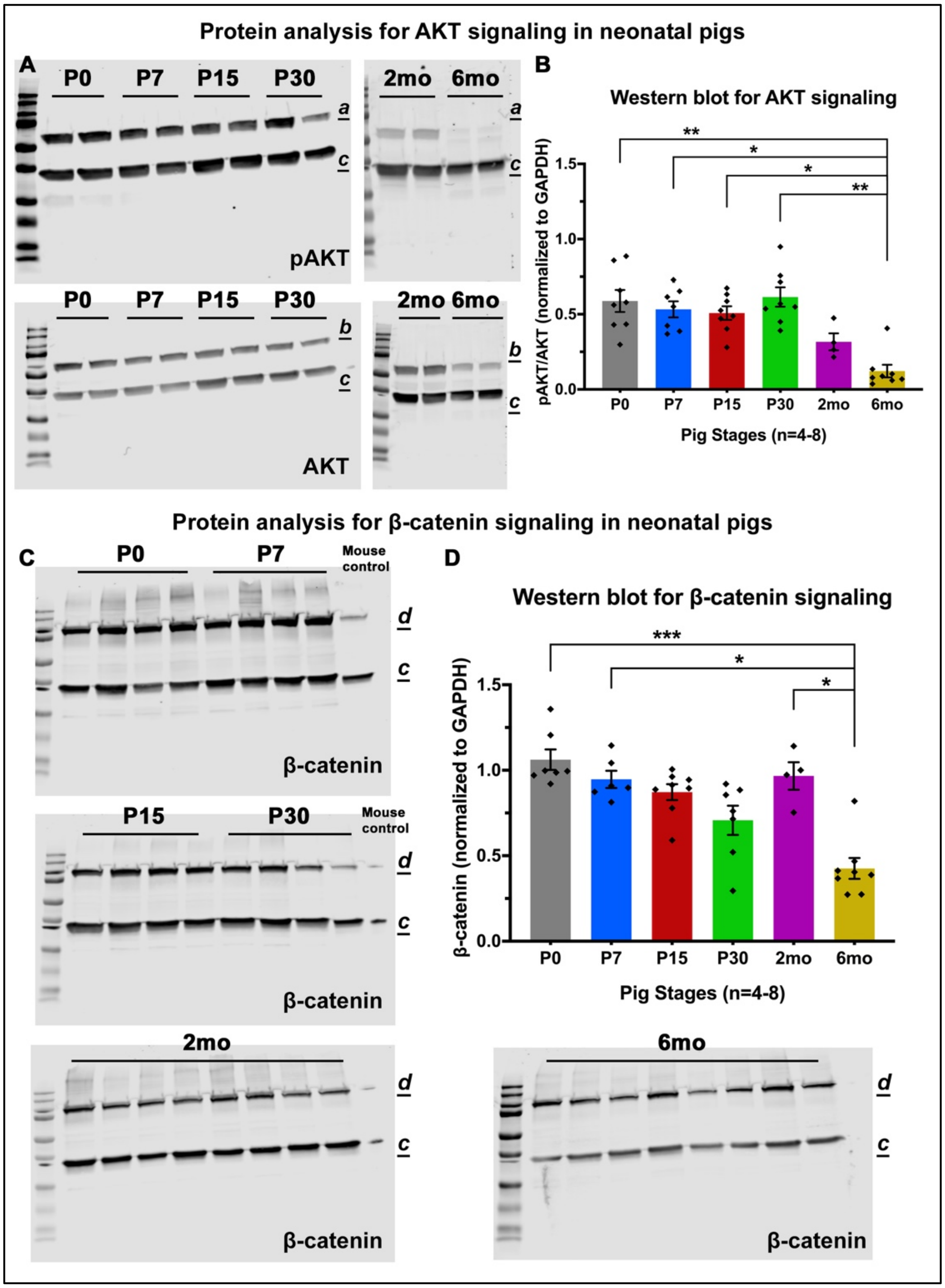


**Figure S6. Cardiac developmental and proliferative signaling is not downregulated until 6mo in pig left ventricles. (A)** Representative Western blots for pAKT and AKT levels in pig left-ventricular tissue at P0-6mo. a indicates pAKT (60kDa), b indicates AKT (60kDa), with c GAPDH (30-40kDa) utilized for normalization. **(B)** Ratios of pAKT/AKT assessed from Western quantification normalized to GAPDH. **(C)** Representative Western blots for β-catenin levels in pig left-ventricular tissue, with P30 mouse ventricular protein as control. d indicates β-catenin (100kDa), with c GAPDH (30-40kDa) utilized for normalization. **(D)** β-catenin levels assessed by Western quantification normalized to GAPDH. Data are mean ± SEM, with *p<0.05, **p<0.01, ***p<0.001 determined by Dunn's Kruskal-Wallis Multiple Comparisons Tests (n=4-8 per stage).

#### Supplementary Tables

**Supplementary Table 1.** Antibodies and reagents utilized in this study.

| Figure No. | Supp. Fig. No. | Antibodies / Reagents | Catalog No. / Supplier |
|---|---|---|---|
| 1 | S1 | Hematoxylin and Eosin (H&E) Staining Kit | American Master Tech Scientific, McKinney, TX |
| 1, 3-7 | S1, 3, 4 | Wheat Germ Agglutinin (WGA) 647 [1:250] | W32466 Thermo Fisher Scientific, Waltham, MA |
| 1-5, 7 | S1, 3, 4 | 4′,6-diamidino-2-phenylindole (DAPI) [1:10000] | D1306 Thermo Fisher Scientific, Waltham, MA |
| 2 | S2 | Lectin (Biotinylated PNA) [1:300] | BA-0074 Vector Laboratories, Burlingame, CA |
| 2 | S2 | Avidin-Biotin (ABC) Kit | PA-6100 Vector Laboratories, Burlingame, CA |
| 2 | S2 | Metal Enhanced DAB Substrate Kit [1X working] | 34065 Thermo Fisher Scientific, Waltham, MA |
| 2 | - | Masson’s Trichrome 2000 Stain Kit | American Master Tech Scientific, McKinney, TX |
| 2 | S3 | Collagen Hybridizing Peptide (CHP) [20μM working] | FLU300 3Helix, Salt Lake City, UT |
| 2, 3, 7 | S3-5 | NucleoSpin RNA Isolation Kit | 40955 Macherey-Nagel, Duren, Germany |
| 2, 3, 7 | S3-5 | DNase I enzyme | EN0255 Thermo Fisher Scientific, Waltham, MA |

| | | | |
|---|---|---|---|
| 2, 3, 7 | S3-5 | cDNA Synthesis Kit (SuperScript III First-Strand Synthesis SuperMix) | 11752250 Thermo Fisher Scientific, Waltham, MA |
| 2, 3, 7 | S3-5 | RiboLock RNase Inhibitor | EO0381 Thermo Fisher Scientific, Waltham, MA |
| 2, 3, 7 | S3-5 | SYBR Green PCR Master Mix | 4367660 Thermo Fisher Scientific, Waltham, MA |
| 3 | S4 | Connexin-43 (Cx43) [1:300] | ab11370 Abcam, Cambridge, UK |
| 3, 6 | S4 | Desmin [1:200] | ab80503 Abcam, Cambridge, UK |
| 4, 5, 7 | - | Sarcomeric $\alpha$-actinin [1:100] | A7811 Millipore Sigma, Burlington, MA |
| 4, 5, 6 | - | Collagenase type 2 [4mg/ml working] | LS004174 Worthington Biochemicals, Lakewood, NJ |
| 4, 5, 6 | - | Collagenase type 4 [1mg/ml working] | LS004186 Worthington Biochemicals, Lakewood, NJ |
| 4, 5, 6 | - | Dispase II [2mg/ml working] | 17105041 Thermo Fisher Scientific, Waltham, MA |
| 6 | - | Hoechst 33342 [1:10000] | H3570 Thermo Fisher Scientific, Waltham, MA |
| 6 | - | Vimentin [1:200] | ab45939 Abcam, Cambridge, UK |
| 7 | - | Phosphohistone H3 (pHH3) [1:100] | 06-570 Millipore Sigma, Burlington, MA |

| | | | |
|---|---|---|---|
| 7 | - | PCM1<br>[1:200] | HPA023370<br>Sigma-Aldrich, St. Louis, MO |
| 7 | - | Wheat Germ Agglutinin (WGA) TRITC<br>[1:250] | W849<br>Thermo Fisher Scientific, Waltham, MA |
| - | S6 | CelLytic MT Cell Lysis Reagent | C3228<br>Sigma-Aldrich, St. Louis, MO |
| - | S6 | Halt Protease and phosphatase inhibitor cocktail | 78440<br>Thermo Fisher Scientific, Waltham, MA |
| - | S6 | Laemmli sample buffer | 1610737<br>Bio-Rad, Hercules, CA |
| - | S6 | Tris/Glycine/SDS running buffer | 1610732<br>Bio-Rad, Hercules, CA |
| - | S6 | Odyssey blocking buffer (TBS) | 927-50000<br>Licor, Lincoln, NE |
| - | S6 | AKT<br>[1:1000] | 9272s<br>Cell Signaling Technology, Danvers, MA |
| - | S6 | pAKT<br>[1:1000] | 4058s<br>Cell Signaling Technology, Danvers, MA |
| - | S6 | β-catenin<br>[1:250] | 71-2700<br>Invitrogen, Carlsbad, CA |
| - | S6 | GAPDH<br>[1:50000] | 10R-G109A<br>Fitzgerald Industries, Tompkinsville, KY |
| - | S6 | Precision Plus Protein™ Dual Color Standards (10-250kDa) | 161-0374<br>Bio-Rad, Hercules, CA |

| | | |
|---|---|---|
| **Common Reagents** | Fish Skin Gelatin Blocking | NC0382999<br>Biotium Inc., Fremont, CA |
| | Citrate Antigen Retrieval Buffer (pH 6.0) | ab93678<br>Vector Labs, Burlingame, CA |
| | Goat Serum | G9023<br>Sigma-Aldrich, St. Louis, MO |
| | Donkey Serum | G9663<br>Sigma-Aldrich, St. Louis, MO |
| | Donkey IgG secondary antibodies [1:500] | Abcam, Cambridge, UK |
| | Goat IgG secondary antibodies [1:500] | Abcam, Cambridge, UK |
| | Hydrogen Peroxide | H1009<br>Millipore Sigma, Burlington, MA |
| | Paraformaldehyde (PFA) | Electron Microscopy Sciences, Hatfield, PA |
| | Formaldehyde | Electron Microscopy Sciences, Hatfield, PA |
| | Xylene | X5-1<br>Fisher Scientific, Hampton, NH |
| | Ethanol | 2701, 2801<br>Decon Labs, PA |
| | TissuePrep2 Embedding Media (Paraffin) | 8002-74-2<br>Fisher Scientific, Hampton, NH |
| | Optimal Cutting Temperature (OCT) Compound | 4585<br>Fisher Scientific, Hampton, NH |
| | Vectashield Hardset Mounting Medium | H-1400<br>Vector Labs, Burlingame, CA |

**<u>Supplementary Table 2</u>.** Primer sequences used for mRNA analysis by RT-qPCR with SYBR Green.

##### 1A. PIG PRIMERS

| ***Gene Name (Pig)*** | **5' - Forward Primer Sequence - 3'<br>5' - Reverse Primer Sequence - 3'** |
|---|---|
| *18S* | AATTCCGATAACGAACGAGACT<br>GGACATCTAAGGGCATCACAG |
| *CCNA2* | CCCTGCATTTGGCTGTGAAC<br>ATTCAGGCCAGCTTTGTCCC |
| *CCNB1* | CATGCAGGATAATTGTGTGCCC<br>CCTCGATTCACCACGACGAT |
| *CCND1* | GCGAGGAACAGAAGTGCG<br>TGGAGTTGTCGGTGTAGATGC |
| *CCNE2* | TCAAGACGCAGTAGCCGTTT<br>AGCCAAACATCCTGTGAGCA |
| *CDK1* | GGAAACCAGGAAGCCTAGCA<br>ACAACGTGTGGGAAAGCTACA |
| *CDK2* | AAGTGGGCCAGGCAAGATT<br>GAGCTGCCTTTGCTGAAATCC |
| *CDK4* | ATGTGGAGCGTTGGCTGTAT<br>TGCTCCAGACTCCTCCATCT |
| *CDKN1A* | ACCATGTGGACCTGTTGCTGT<br>AGAAATCTGTCATGCTGGTCTGCC |
| *CDKN1C* | GGTTATGCCAAAGGCACGTC<br>GACTGCAAGCTAGATGGGCT |
| *CDKN2B* | ACCGTGCGTCAACTTCTGG<br>CAGAAACCGGGCAACGTCA |

| | |
|---|---|
| *CDKN2C* | ACCGAACTGGTTTCGCTGTC<br>CTTTGCTGGCGGTATGCTTT |
| *PRC1* | GTCAAGCATGGAGCCAATGAG<br>AGAATAGGTGCTGGCAACAGA |
| *COL1A1* | CTGGAAGAGCGGAGAATACTG<br>CTGTAGGTGAAGCGGCTGTT |
| *COL3A1* | CCTGGACGAGATGGAAACCC<br>GGCTACCTACTGCACCTTGG |
| *COL4A1* | TTGGCGGTTCTCCAGGAATC<br>GTCACACCCTGCTGTCCTTT |
| *MYH6* | GTGAAGAGATAACCAGAGGAGCG<br>CACCTGATCCTCCTTCACGG |
| *MYH7* | AAGGTCAAGGCCTACAAGCG<br>CTTTGTTGCGCCCTCAGGAT |
| *TNNI1* | AACTTCACGCCAAGGTGGAG<br>ATGGCCTCGACGTTCTTTCT |
| *TNNI3* | ATACGACGTGGAGGCGAAAG<br>CATCATGGCATCGGCAGAGA |
| *SLC2A1* | ATGCGGGAGAAGAAGGTCAC<br>CACGAACAGCGACACGACA |
| *HK1* | ATGTGCGTTTCCTCCTCTCG<br>AAATCCATCTCGGCTCGCAT |
| *LDHA* | GCCCGGTTCCGTTACCTAAT<br>CACCACCTGTTTGTGAACCG |
| *SLC27A1* | ACCTATCAGGTGACGTGCTG<br>ACTCTGATCCAGAGGCAGGT |
| *CPT1B* | AAGTCCTTCACCCTCATCGC<br>TGCCAGCATAGGGTTTGGTT |

| | |
|---|---|
| *ACADM* | ATAGAACTGGCGAGTACCCTG<br>AGGCACACATCAATGGCTCC |
| *HADHA* | ACGAGCTTTGGCTTTCCTGT<br>GCGGTACTGGATGTCTTCGT |

**1B. MOUSE PRIMERS**

| ***Gene Name (Mouse)*** | **5' – Forward Primer Sequence – 3'<br>5' – Reverse Primer Sequence – 3'** |
|---|---|
| *18s* | TTTCTCGATTCCGTGGGTGG<br>TCAATCTCGGGTGGCTGAAC |
| *Ccna2* | CTTGGCTGCACCAACAGTAA<br>ATGACTCAGGCCAGCTCTGT |
| *Ccnb1* | AGCAAATATGAGGAGATGTACC<br>CGACTTTAGATGCTCTACGGA |
| *Ccnd1* | AGTGCGTGCAGAAGGAGATT<br>CACAACTTCTCGGCAGTCAA |
| *Ccne2* | ATGTCAAGACGCAGCCGTTTA<br>GCTGATTCCTCCAGACAGTACA |
| *Cdk1* | GTCCGTCGTAACCTGTTGAG<br>TGACTATATTTGGATGTCGAAG |
| *Cdk2* | TTTGCTGAAATGGTGACCCG<br>GGCTGAAATCCGCTTGTTGG |
| *Cdk4* | CGTGAGGTGGCCTTGTTAAG<br>GTACCAGAGCGTAACCACCA |
| *Cdkn1a* | CGAGAACGGTGGAACTTTGAC<br>CCAGGGCTCAGGTAGACCTT |

| | |
|---|---|
| *Cdkn1c* | TCAGCCAGCCTTCGACCAT<br>TTGAAGTCCCAGCGGTTCTG |
| *Cdkn2b* | CCTTTCAGGACGCGGTGTAA<br>CTGACTGCACCCACCCAAAT |
| *Cdkn2c* | ACCATCCCAGTCCTTCTGTCA<br>AGAAGCCTCCTGGCAATCTC |
| *Prc1* | AAGGAGCTGAGTACCCTGTG<br>CAGAGGATGTCACGGAGTTCT |
| *Col1a1* | CTGGCCTCCCTGGAATGAAG<br>GCTTCACCCTTAGCACCAACT |
| *Col3a1* | GGACACAGAGGCTTTGATGGA<br>CCACCAGGACTGCCGTTATT |
| *Col4a1* | TCATTAGCAGGTGTGCGGTT<br>GCAGAGGCGAGCATCATAGT |
| *Myh6* | CCTCAAGCTCATGGCTACAC<br>GCTGGGTTCAGGATGCGATA |
| *Myh7* | TTACTTGCTACCCTCAGGTGG<br>CAGTCACCGTCTTGCCATTCT |
| *Tnni1* | TTCAGGACTTGTGCCGAGAG<br>TACAGCAAGCCAACCTCTACTG |
| *Tnni3* | TCTGCCAACTACCGAGCCTAT<br>CTCTTCTGCCTCTCGTTCCAT |
| *Slc2a1* | CTCTGTCGGCCTCTTTGTTAAT<br>CCAGTTTGGAGAAGCCCATAAG |
| *Hk1* | TTCGAGAAGATGGTGAGCGG<br>AGAGTTCCCATCCCGTTTCA |
| *Ldha* | GTCCAGCGAAACGTGAACAT<br>TTCCACTGCTCCTTGTCTGC |

| | |
|---|---|
| *Slc27a1* | CGCTTTCTGCGTATCGTCTG<br>GATGCACGGGATCGTGTCT |
| *Cpt1b* | CCTGGTGCTCAAGTCATGGT<br>TCCAGTTTGCGGCGATACAT |
| *Acadm* | ATGCCTGTGATTCTTGCTGGA<br>ACATCTTCTGGCCGTTGATAAC |
| *Hadha* | TGCATTTGCCGCAGCTTTAC<br>GTTGGCCCAGATTTCGTTCA |

### Supplementary Methods

This is an expanded section for the methods provided in the main article. Information on antibodies and reagents is provided in Supplementary Table 1, while primer sequences used for RT-qPCR are listed in Supplementary Table 2.

#### 1 Animals

All experiments involving animals were performed conforming to the NIH Guide for the Care and Use of Laboratory Animals and all protocols involving animals were approved by the Cincinnati Children's Hospital Institutional Animal Care and Use Committee (IACUC).

##### *1.1 Pigs*

A total of n=68 White Yorkshire-Landrace farm pigs were utilized in this study, at ages P0 (postnatal day 0), P7, P15, P30, 2mo (2 months post-birth) and 6mo. The distribution of pigs was as follows: n=14 per stage at P0 to P30, n=4 at 2mo, and n=8 at 6mo. P0 to P30 pigs were purchased from Michael Fanning Farms, (Howe, IN), 2mo pigs from Isler Genetics (Prospect, OH), and 6mo pig hearts were obtained from a local abattoir. Pig euthanasia and cardiac tissue harvests were conducted off-site for P0 to P30 and 6mo pigs, with excised heart tissue pieces prepared and transported to the laboratory at CCHMC (Cincinnati, OH). 2mo pigs were transported to in-house vivarium 2 days before harvest for acclimatization, and subsequently euthanized for heart harvests. Male and female pigs were analyzed at all stages. Only left ventricular free wall tissue was utilized for study from all pigs, with tissue harvested from the same approximate area of each heart (middle region of the left ventricle), to avoid apex-to-base variations in study results. Out of n=14 pigs per stage at P0 to P30, ventricular tissue of n=6 per stage were processed for cardiomyocyte (CM) dissociations (detailed in Section 6), while the other n=8 per stage were used for tissue staining, and mRNA/protein isolation. Ventricular tissue from all pigs at 2mo and 6mo stages was used for tissue staining, mRNA/protein isolation, and CM dissociation.

##### *1.2 Heart and body weights in pigs*

Body weights were obtained by weighing whole pigs (in kg) prior to removal of the heart on a farm animal weighing scale. Following harvest, exsanguinated total hearts were weighed (in g). Heart weight-to-body weight ratios were calculated on Microsoft Excel and visualized using GraphPad Prism 8. Fold-increases in heart and body weights after birth were calculated at P7-6mo relative to heart and body weights at birth (P0).

##### *1.3 Mice*

A total of n=48 wildtype FVB/N mice were bred in-house at the vivarium in CCHMC, Cincinnati, OH (founder animals were originally sourced from Jackson Labs, Bar Harbor, ME), and hearts were harvested at P0, P7, P15, and P30 (n=12 hearts per stage). Hearts were processed for tissue staining (n=3 hearts per stage for paraffin sections, n=3 hearts per stage for cryosections) and mRNA analysis (n=6 hearts per stage). Male and female mice were represented at all stages.

#### 2 Tissue processing

Fresh left ventricular tissue samples from pigs after harvest were washed in 1X phosphate buffered saline (PBS) and subsequently placed in 4% paraformaldehyde (PFA) for tissue staining studies, in 3.7% formaldehyde for CM dissociations, or flash-frozen in liquid nitrogen for mRNA and protein analysis. Fixed hearts in 4% PFA were subsequently embedded in Optimal Cutting Temperature (OCT) compound or dehydrated and embedded in Paraffin wax for cryo- and paraffin-sectioning respectively, as described previously.[1] Microtome sections of 5-7μm were obtained, encompassing the epicardium to endocardium in each tissue section per slide. In mice, whole hearts were washed in 1X PBS prior to fixation in 4% PFA, and whole ventricular tissue was flash-frozen in liquid nitrogen for all stages.

#### 3 Histochemical and immunohistochemical staining in tissue sections

Paraffin sections were dewaxed by Xylene washes followed by rehydration with graded ethanol concentrations, as described previously.[1] Brightfield images were obtained using an Olympus BX51 microscope equipped with a Nikon DS-Ri1 camera (Tokyo, Japan).

##### *3.1 H&E and Trichrome staining*

Hematoxylin and Eosin (H&E) staining kit was utilized according to the manufacturer's protocol provided, to visualize cardiac morphology. Masson's Trichrome 2000 stain kit was used according to manufacturer's protocol to visualize total collagen in tissue sections. Pigs from P0 to 6mo at n=4 per stage were utilized for histological staining and analysis. Mice from P0 to P30 at n=3 per stage were utilized and processed alongside pig samples.

##### *3.2 Lectin-DAB staining*

Antigen retrieval with citrate buffer (1X, pH 6.0) was carried out using pressure cooker/microwave. Blocking was performed with 0.3% hydrogen peroxide and 6% goat serum for 30 minutes each, followed by overnight incubation with biotinylated lectin at 4°C. Staining was detected by Avidin-Biotin Complexing (ABC) kit following manufacturer protocols, followed by color development by immersion for 1 minute in freshly-prepared 3,3'-diaminobenzidine (DAB) metal concentrate in stable peroxidase. Capillaries were identified as small circular Lectin-DAB positive structures. Vascular density was estimated by counting individual Lectin-DAB positive small capillaries in 20X images of at least 4 random cardiac regions per tissue section (n=4-8 pigs per stage), in two technical replicates. Area per 20X image was measured. Both measurements were performed using Fiji (ImageJ) analysis software. Average number of Lectin-DAB positive capillaries per area ($0.3mm^2$) per pig was assessed for vascular density. In mice, the same protocol as above was followed and tissue slides were processed alongside pig samples. At least 3 random 20X images per slide were measured in n=3 mice per stage.

#### 4 Immunofluorescence staining in tissue sections

Confocal images were captured using a Nikon Eclipse Ti Fluorescence microscope, with NIS elements software (Tokyo, Japan). Paraffin-embedded tissue sections were deparaffinized as described above (Section 3). Cryosections were thawed at room temperature followed by 1X PBS washes. Unless specified otherwise, 1X citrate buffer antigen retrieval was performed using pressure cooker/microwave for 20 minutes (paraffin sections) or by incubation of slides for 1 hour at room temperature (cryosections). Unless specified otherwise, blocking was performed using 1X Fish skin gelatin blocking buffer for 1-2 hours at room temperature.

##### *4.1 Cardiac morphology and ventricular nuclear density assessment*

Paraffin sections following deparaffinization and antigen retrieval were stained with Wheat Germ Agglutinin Alexa Fluor 647 conjugate (WGA 647, cell membrane) and DAPI (nuclei), in order to visualize cardiac morphology in pigs (n=4 per stage) and mice (n=3 per stage). Approximate ventricular nuclear density (number of nuclei per area) per stage in pigs was measured by Object Counts for individual nuclei via DAPI (blue) pixel identification in NIS Elements software, in 60X images of 4 random cross-sectional cardiac regions per tissue section (n=4 per pigs per stage). Average number of nuclei per area (0.05mm$^2$) per pig was graphed using GraphPad Prism 8.

*4.2 Collagen hybridizing peptide staining*

Cryosections were rehydrated by 1X PBS washes. Collagen Hybridizing Peptide (CHP) working solution (20μM) was freshly prepared by heating to 80°C then cooling in ice for 10-15 seconds. Tissue samples were incubated in CHP working solution overnight at 4°C. CHP conjugated to 5-FAM was visualized with excitation/emission at 495/520nm. Co-staining with DAPI was performed following CHP incubation to visualize nuclei. 60X images of at least 4 random cardiac regions per tissue section (n=4-8 pigs per stage), in two technical replicates, were obtained in pigs. CHP area was measured by fluorescence thresholding in NIS elements. Total area per image was measured in NIS elements. The ratio of CHP-stained area to total measured area (0.05mm$^2$) per image was calculated to obtain the percentage of remodeling collagen per area. In mice, n=3 cryosections per stage were processed alongside the pig samples.

*4.3 Connexin-43 staining for sarcomeric gap junctional maturation*

Gap junctional maturation was identified by anti-Connexin-43 (Cx43, a gap junctional protein) staining in cryosections, with anti-Desmin to visualize CM. Thawed and washed cryosections were permeabilized in 0.1% Triton for 15 minutes, followed by blocking in 3% BSA for 1-2 hours. Primary antibody was incubated overnight at 4°C, followed by Donkey IgG secondary antibodies, WGA 647 (cell membrane) and DAPI (nuclei) co-staining. 60X images were obtained in n=4 pigs per stage and n=3 mice per stage to visually assess localization of Cx43 to CM termini identified by WGA membrane staining.

*4.4 Cardiomyocyte cross-sectional area and nuclear area assessment*

Paraffin sections following deparaffinization and antigen retrieval were stained with WGA 647 (cell membrane) and DAPI (nuclei) to assess CM cross-sectional area (CSA, μm$^2$). Fiji (ImageJ) analysis software was used to measure CSA by individual cell tracing in 20X images of 4 random cardiac regions per paraffin tissue section (n=4-8 pigs per stage), and in two technical replicates. In the same images, area of individual nuclei was determined by Object Area (μm$^2$) measurements with DAPI staining in NIS Elements software.

*4.5 Phosphohistone-H3 staining for cardiomyocyte mitotic activity*

Cryosections following citrate antigen retrieval and blocking were stained with anti-Phosphohistone-H3 Ser10 (pHH3) antibody, in combination with anti-Sarcomeric α-actinin to identify CMs, WGA 647 or WGA TRITC to identify cell membranes, and DAPI staining of nuclei. Additional cryosections were stained with PCM1 to identify CM nuclei with WGA 647, and DAPI. Primary antibodies were incubated overnight at 4°C. Goat IgG secondary antibodies were utilized. In 20X images of 4-6 random cardiac regions per tissue section (n=4-8 pigs per stage), and in two technical replicates, pHH3+ CM nuclei as identified by α-actinin or PCM1 were counted manually in NIS Elements software. The total number of nuclei per image was counted by Object Counts of DAPI staining in NIS Elements. Mitotic indices were calculated by ratios of pHH3+ CM nuclei to total CM nuclei and the results visualized on GraphPad Prism 8.

#### 5 mRNA analysis by RT-qPCR

Ventricular mRNA from n=4-8 pigs per stage and n=6 mice per stage was isolated for gene expression analysis by quantitative real-time polymerase chain reaction (RT-qPCR). Flash-frozen tissue samples from pig and mouse ventricles harvested as described above (Section 2) were stored at -80°C until needed. RNA was extracted using NucleoSpin RNA kit following manufacturer's protocol. Further purification was carried out by treatment with DNaseI/RNase inhibitor cocktail followed by cDNA synthesis using SuperScript III First-Strand Synthesis kit, per manufacturer's protocols. RT-qPCR was performed using SYBR Green. Gene-specific primer sets for pigs and mice (Supplementary Table 2) were designed using NCBI PrimerBlast and validated by Sanger sequencing. Samples were amplified in duplicates for 35 cycles using StepOnePlus Real-Time PCR machine (Applied Biosystems). Relative gene expression was calculated by comparative $\Delta\Delta Ct$ method.[2] *18S* rRNA expression was used for normalization. Normalized average expression of P0 was set to 1.0 to calculate fold change after birth in mice and pigs.

#### 6 Cardiomyocyte dissociations

The protocol of Mollova *et al.*[3] for CM cell dissociation from formalin-fixed tissue was modified for isolating CMs from pig ventricular tissue. Pig heart pieces of about 2mm in length were excised from ventricular tissue fixed in 3.7% formaldehyde overnight at 4°C, or from tissue stored in 10% neutral-buffered formalin at 4°C. Heart pieces were washed in 1X PBS and subsequently digested by rocking at 37°C for 48 hours in freshly-prepared digestion mixture (4mg/ml Collagenase type 2, 1mg/ml Collagenase type 4, and 2mg/ml Dispase II in 1X PBS). After digestion, eluate was filtered through 200-micron nylon mesh (NC0148096, Fisher) and gently centrifuged at 800rpm for 2 minutes to obtain CM pellets. Resuspended pellets were utilized for antibody staining. Staining steps were performed by repeated centrifugations as described above, followed by pellet resuspensions in appropriate solutions. Following antibody staining, final cell suspensions were pelleted, resuspended in a few drops of Vectashield Hardset mounting medium, deposited onto slides using coverslips, and dried overnight at 4°C before imaging. Individual whole CMs were visually identified based on sarcomeric staining.

##### *6.1 Nucleation counts in cardiomyocytes*

Dissociated CMs were stained with anti-Sarcomeric α-actinin overnight at 4°C, followed by Goat IgG secondary antibody staining and DAPI. Stained CMs were deposited on slides as described above and confocal images at 20X with at least 10 images per slide and in two technical replicates (n=4-5 pigs per stage) were obtained for assessment of nucleation. The number of nuclei per CM was counted manually in NIS Elements software. Percentages of mono-, bi-, and multinucleated CMs out of total number of CMs counted per pig in each stage was calculated and the results plotted using GraphPad Prism 8.

##### *6.2 Cardiomyocyte total surface area, length, and width measurements*

CM dissociations were stained with anti-Sarcomeric α-actinin and DAPI as described above. Total surface area per CM ($\mu m^2$) was measured after manually tracing cell outlines in at least 6 random 20X images per slide (n=3-5 pigs per stage) and in two technical replicates, using Fiji (ImageJ) software. In the same images, CM lengths and widths (µm) were individually measured by manual tracing of longest (end-end) and widest (middle) points respectively, using Fiji (ImageJ) software. Averages of measurements per pig per stage were calculated and plotted using GraphPad Prism 8.

**7 Assessment of nuclear DNA content in dissociated cardiomyocytes and tissue sections**

The method of Patterson *et al.*[4] was modified for determination of nuclear DNA content in pig CMs. Pig CM dissociations prepared as described above were mixed with formalin-fixed adult male farm pig sperm after PBS washes. Dissociated cells were stained with anti-Desmin to identify CMs and Hoechst 33342 for nuclear DNA quantification. Confocal images at 60X were acquired with at least 4 images per slide, in n=4 pigs per stage. Hoechst intensities of each nucleus per image were measured by Hoechst 33342 (blue stain) thresholding in NIS Elements software. CM versus sperm nuclei were identified manually in each image by Binary Object counting. CM nuclear intensities were then normalized to sperm nuclear intensities (which are set to 1.0), to obtain relative intensity values. The normalized relative intensity values of individual CM nuclei were plotted using GraphPad Prism 8. For assessment of nuclear DNA content in cryosections, antigen retrieval and blocking were performed as described above, and sections were stained with Desmin, Hoechst 33342, Vimentin (non-CM marker), and WGA 647. Primary antibodies were incubated overnight at 4°C, and Donkey IgG secondary antibodies were utilized. Images were acquired at 60X with at least 4 images per slide (n=4 pigs per stage), and Hoechst intensity of each nucleus per image was measured in NIS Elements as described above. Desmin-positive (CM) cell nuclei versus Vimentin-positive (non-CM) cell nuclei and their corresponding nuclear intensities were identified manually by Binary Object counting. Hoechst intensity of each CM and non-CM nucleus per image was then normalized to the average of non-CM (diploid, 2c) nuclear intensities for that image, to obtain normalized relative intensity values. Normalized relative nuclear intensity values close to 1.0 in this analysis indicate 2c DNA content. The normalized relative intensity values of individual CM and non-CM nuclei were plotted using GraphPad Prism 8. Based on standard deviation of non-CM relative nuclear intensity values from P0-6mo, thresholds were allotted as: below 1.3 (2c) and above 1.3 (>2c), no units. In longitudinal sections, multinucleated CMs were identified to similarly measure individual nuclear intensities within each multinucleated CM. Utilizing the above thresholds, the percent of nuclei with normalized relative Hoechst intensity values at 2c (below 1.3) or >2c (above 1.3) was calculated for each pig per stage, and the results presented using GraphPad Prism 8.

**8 Western blotting for AKT and β-catenin signaling in pig ventricular protein**

Pig ventricular tissue stored in -80°C was utilized. Protein was extracted from pig myocardial tissue by homogenizing in CelLytic MT cell lysis reagent at a ratio of tissue to reagent at 1:20. Phosphatase inhibitor cocktail and protease inhibitor cocktail were added to the cell lysis reagent, followed by centrifugation and collection of protein supernatant. Protein quantification was performed using pre-set BSA program on Direct Detect software (Merck Millipore). Samples (1.5µg protein/µl) were prepared in Laemmli sample buffer and Cell lysis reagent. Following denaturation at 85°C for 15 minutes, 20µl was run on a 12% Mini-PROTEAN TGX precast gel (Bio-Rad) in Tris/Glycine/SDS running buffer with Precision Plus Protein molecular weight ladder (10–250kDa). Following protein transfer to an Immobilon-FL transfer membrane (Merck Millipore), non-specific binding was blocked by incubation in TBS blocking buffer for 1 hour. Membranes were incubated with primary antibodies: AKT, pAKT, β-catenin, and GAPDH overnight at 4°C. Membranes were then washed and incubated with fluorophore-conjugated secondary antibodies followed by quantification of AKT, pAKT or β-catenin protein levels, relative to GAPDH, using an Odyssey CLx infrared imaging system (Licor). Normalized expression values were plotted as pAKT/AKT and β-catenin levels for n=4-8 pigs per stage.

**9 Graphical representation and statistics**

Statistical analysis and graph preparation were done using GraphPad Prism 8 software. Numerical data are shown as means ± SEM, unless individual sample distribution is not provided in the graph in which case means ± SD is displayed (as noted appropriately in figure legends). Statistical significance was determined by unpaired One-way ANOVA non-parametric Multiple Comparisons tests such as Kruskal-Wallis test with Dunn's corrections or Brown-Forsythe and Welch test with Games-Howell corrections. $p<0.05$ (*) was deemed significant, with $p<0.01$ (**), $p<0.001$ (***), and $p<0.0001$ (****) also represented.
